# Interaction of modified oligonucleotides with nuclear proteins, formation of novel nuclear structures and sequence-independent effects on RNA processing

**DOI:** 10.1101/446773

**Authors:** Loren L Flynn, Ruohan Li, May T Aung-Htut, Ianthe L Pitout, Jack A L Cooper, Alysia Hubbard, Lisa Griffiths, Charlie Bond, Steve D Wilton, Archa H Fox, Sue Fletcher

## Abstract

Oligonucleotides and nucleic acid analogues that alter gene expression are showing therapeutic promise for selected human diseases. The modification of synthetic nucleic acids to protect against nuclease degradation and to influence drug function is common practice, however, such modifications may also confer unexpected physicochemical and biological properties. Here we report backbone-specific effects of modified oligonucleotides on subnuclear organelles, altered distribution of nuclear proteins, the appearance of novel structured nuclear inclusions, and modification of RNA processing in cultured cells transfected with antisense oligonucleotides on a phosphorothioate backbone. Phosphodiester and phosphorodiamidate morpholino oligomers elicited no such consequences. Disruption of subnuclear structures and proteins elicit severe phenotypic disturbances, revealed by transcriptomic analysis of fibroblasts exhibiting such disruption. These data suggest that the toxic effects and adverse events reported after clinical evaluation of phosphorothioate nucleic acid drugs may be mediated, at least in part, by non-specific interaction of nuclear components with the phosphorothioate backbone.

## Introduction

Antisense oligonucleotide drugs are a class of therapeutics designed to alter gene expression and function, and are reported to have delivered therapeutic benefit to spinal muscular atrophy (SMA) type 1 patients (Paton 2017) and a subset of Duchenne muscular dystrophy (DMD) cases (Mendell *et al.* 2013, Mendell *et al.* 2016). *Nusinersen* (*Spinraza*) is a 2′ O-methoxyethyl antisense oligonucleotide (AO) on a phosphorothioate backbone, targeting a splice silencer (*ISS-N1*) in *SMN2* intron 7 that promotes exon 7 selection during pre-mRNA splicing (Singh *et al.* 2006) and received Food and Drug Administration (USA) (FDA) approval in December 2016 for the treatment of SMA. *Exondys51* is a phosphorodiamidate morpholino oligomer (PMO) that targets the *DMD* pre-mRNA to exclude exon 51, in order to re-frame the dystrophin transcript around frame shifting deletions flanking exon 51, as a treatment for DMD (Mendell *et al.* 2013). *Exondys51* received accelerated approval from the FDA on September 19th 2016. Other drugs exploiting the antisense concept include Kynamro^®^ (mipomersen sodium) (Wong and Goldberg 2014), an adjunctive treatment to reduce LDL-C levels in familial hypercholesteremia, approved in January 2013, followed by an FDA hepatotoxity black box warning; and a number of compounds at various stages of clinical development, predominantly siRNA and oligodeoxyribonucleotide analogues designed to induce RNAse H degradation of target transcripts (for review see (Bennett *et al.* 2017, Stein and Castanotto 2017)). Recently, Alnylam announced the first FDA approval of an RNAi Therapeutic, ONPATTRO™ (patisiran) for the treatment of adult patients with polyneuropathy of hereditary transthyretin-mediated amyloidosis. (https://www.fda.gov/NewsEvents/Newsroom/PressAnnouncements/ucm616518.htm)

Strategies to improve biological stability and confer pharmaceutical properties to nucleic acid drugs include chemical modifications of the bases and nucleic acid backbone to increase resistance to endogenous nucleases; modifications that influence specific oligomer activity and conjugates that enhance cellular uptake and improve tissue distribution. While the phosphorothioate backbone is the most widely applied chemical modification, changes to the ribose moiety (eg, 2’ O-methylation, 2’ O-methoxyethyl, 2’ O-fluoro), nucleobases and other backbone modifications, including peptide nucleic acids and phosphorodiamidate morpholino oligomers can confer specific characteristics and mechanisms of action (for review see (Veedu 2015, Wilton *et al.* 2015, Tri Le *et al.* 2016)).

Many laboratories, including our own, routinely use 2’ O-methyl phosphorothioate oligonucleotides to evaluate antisense sequences for targeted splicing modulation, and different modified bases on a phosphorothioate backbone for various other molecular interventions (Lipi *et al.* 2016, Chakravarthy *et al.* 2017). These compounds can be economically prepared in-house and efficiently transfected into a range of cells as cationic liposomes or dendrimer complexes, using commercially available reagents. However, for sustained splice modification, *in vivo* use and clinical application, PMOs have proved safe and effective (Gebski *et al.* 2003, Fletcher *et al.* 2006, Kinali *et al.* 2009, Mendell *et al.* 2016). We previously reported that antisense exon skipping sequences prepared and optimized *in vitro* as 2’ O-methyl phosphorothioate oligonucleotides generally perform comparably when synthesized as PMO (Adams *et al.* 2007). An extended antisense sequence targeting the *ISS-N1* region of *SMN2* was validated *in vitro* and *in vivo* using 2’ O-methyl phosphorothioate and PMO chemistries (Mitrpant *et al.* 2013).

Phosphorothioate backbone modification of nucleic acid drugs is common practice for protection from rapid degradation by circulating and intracellular nucleases, but these modified AOs can interact undesirably with endogenous cellular components. Early work indicated that negatively charged phosphorothioate AOs bind non-specifically to heparin-like proteins, laminin and collagen (Dias and Stein 2002) and more recently, phosphorothioate AOs were reported to interact with nuclear paraspeckle-associated proteins, sequestering them away from their endogenous target, the long non-coding RNA *NEAT1*, and also seeding paraspeckle like-structures in the absence of *NEAT1* (Shen *et al.* 2014, Shen *et al.* 2015). Paraspeckles and paraspeckle proteins are involved in transcriptional regulation, transport and splicing pathways (Bond and Fox 2009, Naganuma *et al.* 2012). Beyond the paraspeckle-associated subset (Shen *et al.* 2014), phosphorothioate AOs were also reported to bind an additional approximately 50 intracellular proteins (Liang *et al.* 2015) however, these interactions were studied in the context of evaluating the modest impact these interactions had on AO function (Liang *et al.* 2015). A broader question relates to the global consequences on cellular processes arising as a result of the interaction of AOs with nuclear proteins.

Results of *in vivo* studies investigating the consequences of non-specific phosphorothioate AO binding are particularly relevant when considering therapeutic applications. The sequestration of paraspeckle proteins, in particular proteins of the drosophila behaviour/human splicing (DBHS) family, was associated with acute hepatotoxicity, inflammation and apoptosis after application of 2’ fluoro-modified phosphorothioate antisense oligonucleotides with a 5-10-5 gapmer configuration in mice (Shen *et al.* 2018) and while this study focused on the severity of 2’ fluoro-modified phosphorothioate AOs, all 2’ modifications evaluated have been shown to sequester paraspeckle proteins, albeit to varying degrees (Shen *et al.* 2014, Liang *et al.* 2015, Shen *et al.* 2018). The backbone-dependent binding of phosphorothioate-modified olionucleotides to platelets *in vitro* and *in vivo*, mediated by the platelet-specific receptor glycoprotein VI (GPVI) is also of note (Flierl *et al.* 2015), considering the broad range of nucleic acid therapeutics currently under investigation (for review see (Bennett *et al.* 2017). Oligonucleotides on a phosphorothioate-modified backbone elicited strong platelet activation, signaling, reactive oxygen species generation, adhesion, spreading, aggregation and thrombus formation (Flierl *et al.* 2015). Of particular relevance to current clinical usage, 2’ O-methyl phosphorothioate AOs were reported to activate innate immunity when administered directly to the central nervous system (Toonen *et al.* 2018).

Here we report that transfection of cultured cells with 2’ O-methyl phosphorothioate antisense sequences resulted in numerous novel, large nuclear inclusions in the form of highly structured fibril-like aggregates that co-stained for the paraspeckle proteins, NONO, SFPQ, PSPC1 and FUS. Other nuclear proteins showed altered distribution in 2’ O-methyl phosphorothioate transfected cells. Intranuclear inclusions begin to form within four hours of transfection, and become dominant structures throughout the nucleus within 24 hours. The inclusions appear stable once formed and may remain evident on the culture substrate, even after death and disintegration of the cell. Transmission electron microscopy on transfected cells revealed numerous large, regular structures reminiscent of amyloid deposits, with electron dense regions. Furthermore, gene ontology analyses following RNA sequencing demonstrated significant disruptions to chromatin silencing; regulation of autophagy; nucleotide excision repair; membrane and organelle organization; apoptosis; signalling and protein transmembrane transport, following 2’ O-methyl phosphorothioate transfection.

## Materials and Methods

### Antisense oligonucleotides

Phosphorodiamidate morpholino oligomers (PMOs) were purchased from Gene-Tools LLC (Philomath, OR, USA), and 2’ O-methyl phosphorothioate AOs were synthesised by TriLink BioTechnologies (San Diego CA, USA). The following sequences were evaluated; an AO encompassing the *SMN* intron 7 *ISS-N1* target, but with a longer sequence *SMN7D(−10-29)* (5’AUUCACUUUCAUAAUGCUGG 3’); the Gene-Tools standard control oligomer, only known to be biologically active in reticulocytes carrying a splice mutation in the human beta-globin pre-mRNA, as an unrelated sham control AO sequence (5’CCUCUUACCUCAGUUACAAUUUAUA 3’), and identified as ‘control AO’ throughout this study; *Smn* (5’CAC CUU CCU UCU UUU UGA UU 3’) designed to induce exon mouse *Smn* exon skipping and a *SMN* sense AO (5’ CCAGCAUUAUGAAAGUGAAU 3’) complementary to *SMN7D(−10-29).*

### Transfection

The 2′ O-methyl phosphorothioate AOs were transfected into dermal fibroblasts as lipoplexes using 3 μl of Lipofectamine 3000 (Life Technologies, Melbourne, Australia) per 1 ml of OptiMEM, according to the manufacturer’s protocol. The transfection mix was applied to cells seeded at 10,000 per cover slip for immunofluorescence and 15,000 cells per well in 24 well plates for RNA extraction and incubated (37°C) for 24 hours prior to RNA and protein analysis. SMA patient fibroblasts were used for the experiments shown in Figures 1f and 1g (Coriell Cell Repositories, GM03813) and fibroblasts from our in-house biobank, obtained from a healthy volunteer, with informed consent (Murdoch University Human Research Ethics Committee Approval #2013/156) were used for all other experiments, except live-cell imaging (see below).

**Figure 1.**
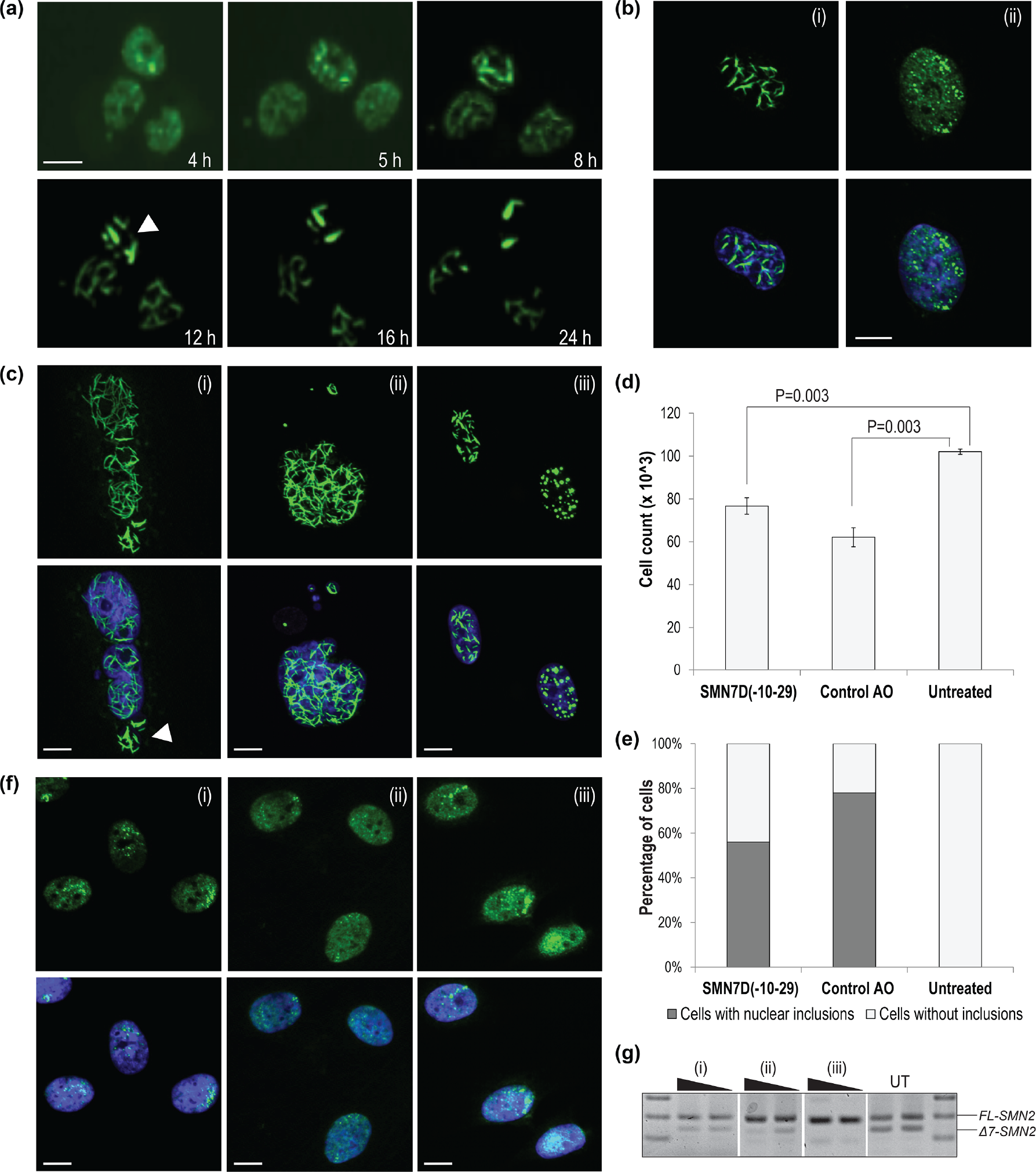
Formation of nuclear inclusions. Showing **(a)** live cell imaging time-course of U2OS cells with endogenous SFPQ-GFP showing the formation of nuclear inclusions over 24 hours following 2’ O-methyl phosphorothioate control AO transfection (100 nM), inclusions are indicated by the white arrow; **(b)** fibroblasts transfected with 12.5 nM of 2’ O-methyl phosphorothioate *SMN7D(−10-29)* stained for NONO (i), compared to untreated fibroblasts (ii); **(c)** fibroblasts stained for NONO show, (i) nuclear inclusions without evidence of nuclear staining after 2’ O-methyl phosphorothioate *SMN7D(−10-29)* transfection, (ii) nuclear blebbing after 2’ O-methyl phosphorothioate control AO transfection and (iii) nuclear inclusions in the form of filaments or foci induced by transfection with a 2’ O-methyl phosphorothioate *SMN7D(−10-29)* sense sequence AO; **(d)** graph showing the number of viable fibroblasts following 2’ O-methyl phosphorothioate AO transfection (n=4), error bars represent standard error of the mean and P-values were calculated using an unpaired T-test; **(e)** graph showing the percentage of fibroblasts containing nuclear inclusions following 2’ O-methyl phosphorothioate transfection with a minimum of 200 cells counted per sample; **(f)** fibroblasts stained for NONO after transfection with PMO *SMN7D(−10-29)* at (i) high dose un-complexed (10 μM), (ii) using a complementary leash and transfection reagent (200 nM) and (iii) using electroporation (1 μM), all incubated for 72 hours; and **(g)** RT-PCR of full length (FL) and exon 7 skipped (Δ7) *SMN2* transcripts of RNA collected following transfections as in **(f)**. All scale bars = 10 μm.

PMOs were delivered to cells either uncomplexed (10 μM), annealed to a complementary sense DNA oligonucleotide (phosphodiester leash) and transfected as a lipoplex (200 nM), or by nucleofection using a NucleofectorTM X Unit (Lonza, Melbourne, Australia) according to the manufacturer’s instructions. PMOs were delivered using the P2 nucleofection kit and CA-137 program at 1 μM, as determined by the final transfection volume. All PMO transfections were incubated for 72 hours prior to RNA and protein analysis.

### RT-PCR on AO transfected cells

RNA was extracted using the MagMAX-96 Total RNA Isolation Kit, including DNase treatment (Life Technologies), according to the manufacturer’s instructions. RT-PCRs were performed using the One-step Superscript III RT-PCR kit with Platinum Taq polymerase (Life Technologies) according to the manufacturer’s instructions. All primer sequences and PCR conditions used in this study are detailed in Supplementary File 1, **Table 1**.

### cDNA synthesis and quantitative PCR

cDNA was synthesised from (~500 ng) RNA extracted from treated and untreated cell cultures using the Superscript IV first-strand synthesis system (Life Technologies) as per the manufacturer’s instructions. Prior to qPCR amplification, cDNA was diluted in RNase/DNase free water (1:5).

The qPCR reactions were performed using Fast SYBR™ Green Master Mix (ThermoFisher Scientific, Melbourne, Australia) and primers for ribosomal RNA subunits *5S* (100% primer efficiency), *18S* (95% primer efficiency), *45S* (100% primer efficiency) and housekeeping Tata box protein (*TBP;*100% primer efficiency) and beta Tubulin (*TUBB;*94% primer efficiency) transcripts. All primer sequences and cycling conditions used in this study are detailed in Supplementary File 1, **Table 1**. Transcript abundance was measured using the CFX384 Touch™ Real Time PCR detection system (Bio-Rad Laboratories Pty., Ltd., Gladesville, Australia). The relative expression of each ribosomal RNA subunit to *TBP* and *TUBB* mRNA was calculated using the 2-DDCT method and presented as a fold change compared to untreated cells.

### Immunofluorescence

Approximately 10,000 fibroblasts were seeded onto 22 mm x 22 mm coverslips in 6-well plates and incubated for 24 hours, prior to transfection. Following transfection, the cells were fixed using acetone: methanol (1:1) on ice for 4 minutes and air-dried. Fixed cells were washed in PBS containing 1% Triton X-100 to permeabilise the nuclear membrane, and then in PBS to remove excess Triton X-100. Primary antibodies were diluted in PBS containing 0.05% Tween20 and applied to cells for 1 hour at room temperature. All antibody details and staining conditions are listed in Supplementary File 1 **Table 2**. Primary antibodies were detected using AlexaFluor488 anti-mouse (cat no. A11001), anti-rabbit (cat. No A11008) or AlexaFluor568 anti-mouse (cat. No A11004) or anti-rabbit (cat. No A11011) (1:400) after incubation for 1 hour at room temperature, and counterstained with Hoechst 33342 (Sigma-Aldrich) for nuclei detection (1 mg/ml, diluted 1:125).

### Western blotting

Cell lysates were prepared with 125 mM Tris/HCl pH 6.8, 15% SDS, 10% Glycerol, 1.25 μM PMSF (Sigma-Aldrich, NSW, Australia and 1x protease inhibitor cocktail (Sigma-Aldrich) and sonicated 6 times (1 second pulses) before adding bromophenol blue (0.004%) and dithiothreitol (2.5 mM). Samples were heated at 94°C for 5 minutes, cooled on ice and centrifuged at 14,000 x g for 2 min before loading onto the gel.

Total protein (10 μg), measured by BCA, was loaded per sample on a NuPAGE Novex 4-12% BIS/Tris gel (Life Technologies) and separated at 200 V for 55 minutes. Proteins were transferred onto a Pall Fluorotrans polyvinylidene fluoride (PVDF) membrane at 350 mA for 2 hours. Following blocking for 1 hour, the membrane was incubated in 5% skim milk powder in 1x TBST containing the primary antibody diluted as shown in Supplementary File 1, **Table 2**. Immunodetection was performed using the Western Breeze Chemiluminescent Immunodetection System (Life Technologies) according to the manufacturer’s instructions. Western blot images were captured on a Vilber Lourmat Fusion FX system using Fusion software and Bio-1D software was used for image analysis.

### Live cell imaging

U2OS cells, previously modified to tag the endogenous SFPQ gene with GFP were used for live-cell imaging (Li *et al.* 2017). Cells were seeded at 1.5 × 105 cells per well in a 12-well plate, 24 hours before transfection, with high glucose DMEM (Life Technologies, Cat No.11995065) supplemented with 10% fetal calf serum and 1% penicillin-streptomycin. On the following day, cells in each well were transfected with 100 nM 2’ O-methyl phosphorothioate AOs complexed with 2.2 μl Lipofectamine 3000 following the manufacturer’s instructions. Immediately after transfection, the plate was transferred to an Incucyte S3 (Essen Bioscience) for live-cell imaging at 1 hour intervals under 10 x optical zoom. SFPQ-GFP and activated caspase-3/7 were visualised under standard green and red channels. Incucyte Caspase-3/7 Red Apoptosis Assay Reagent (Essen Bioscience, Cat No. 4704) was used to reveal cell apoptosis events.

### High resolution microscopy

SIM imaging was performed using an N-SIM microscope (Nikon Corporation, Tokyo, Japan), with SR Apochromat TIRF 100 × 1.49 NA oil immersion objective. Spherical aberration was reduced using the Ti2 automated correction collar at the beginning of the imaging session. Images were acquired using 405 nm, 488 nm and 561 nm lasers, with stacks of step size 0.12 μm (with top and bottom of samples determined visually), using 3D-SIM mode. Images were reconstructed with NIS Elements software (Nikon Corporation, Tokyo, Japan).

### Transmission electron microscopy

Following transfection, cells were washed in PBS and fixed in cold 2.5% phosphate buffered glutaraldehyde overnight. The fixed cells were scraped and centrifuged, then embedded in 4% agarose. The pellets were processed using a Leica tissue processor, moving through 1% aqueous osmium tetroxide, increasing graded alcohols, propylene oxide, propylene oxide/araldite mix, and finally pure araldite resin. The osmicated cells, surrounded by resin in a beem capsule were polymerised overnight at 80oC to form hard blocks. Semi-thin (0.5 micron) sections were taken from the blocks using glass knives, on an RMC ultratome, and stained with methylene blue (0.1% aqueous Methylene Blue with 0.1% Borax) to visualise available cells. Ultrathin sections were then cut from selected blocks, using a Diatome diamond knife, at approximately 95 nm thickness and mounted on copper mesh grids and stained with uranyl acetate (5% uranyl acetate solution in 5% aqueous acetic acid) and Reynold’s lead citrate. The grids were viewed using a JEOL 1400 TEM, at 80 kV and images captured by an 11-megapixel GATAN digital camera at varying magnifications.

### RNA sequencing and analysis

RNA quality was confirmed using a Bioanalyser (Perkin Elmar, MA, USA) prior to RNAseq. Samples were sent to the Australian Genome Research Facility (AGRF, Perth and Melbourne, Australia) for whole transcriptome library preparation using the TruSeq Stranded Total RNA Library Prep Kit (Illumina, CA, USA) and ribosomal RNA depletion with the Ribo-Zero-Gold kit (Illumina, CA, USA). Sequencing was performed using an Illumina HiSeq 2500 (Illumina, CA, USA) to generate 100 base pair single end reads, resulting in an average 25 million reads per sample. Raw sequencing files were quality checked using FastQC (0.11.7), with all files passing. No adapter contamination was identified, and reads were not trimmed. Transcript quantification was performed with salmon (0.8.2), using an index constructed from the Ensembl GRCh38.93 annotation. Transcript abundance was summarised to gene-level counts and imported into R using tximport (1.4.0). Differential expression analysis was performed using DESeq2 (1.16.1) and the default parameters (alpha = 0.1). The heatmap of differentially expressed genes (padj < 0.1) was constructed using the heatmap.2 function from R package gplots (3.0.1). Gene ontology analysis was performed using GSEA (3.0) with 1000 gene set permutations. The gene ontology network was constructed using the Enrichment Map plugin for Cytoscape (3.5.1), with cut-offs: p < 0.05, FDR < 0.1 and edge > 0.3.

## Results

### Formation of nuclear inclusions after 2’ O-methyl phosphorothioate AO transfection

The phosphorothioate backbone can interact non-specifically with both intra- and extra-cellular proteins, and recruits paraspeckle proteins and forms paraspeckle-like structures when transfected into Hela cells (Shen *et al.* 2014). However, the dynamics of the formation of these structures, as well as the long-term consequences of these structures on cell physiology remain unknown.

In this study, we largely focused on two 2’ O-methyl phosphorothioate AOs, one that promotes *SMN2* exon 7 retention during splicing by targeting the *ISS-N1* intronic silencer motif (Singh *et al.* 2006) and the other a control oligomer sequence reported by Genetools LLC and widely used as a transfection control, although additional AO sequences were also tested in some experiments, as indicated below. In order to follow nuclear inclusion formation, we first transfected 2’ O-methyl phosphorothioate AO as lipoplexes (100 nM) into U2OS cells expressing genome engineered GFP-labelled endogenous SFPQ (Li *et al.* 2017) and performed live cell imaging over 48 hours (**Figure 1a**). The AO transfection induced formation of nuclear inclusions within 5 hours, and over a period of 12 hours, all of the SFPQ within the nucleus is observed sequestered within the inclusions, as revealed by decreased SFPQ staining of the nucleoplasm. Ultimately, only the inclusions are evident, at which time, the inclusions then coalesce into discrete filaments.

Immunofluorescent staining of the paraspeckle protein, NONO, showed colocalization in nuclear inclusions in fibroblasts transfected with 2’ O-methyl phosphorothioate *SMN7D(−10-29)* sequence at 100 nM concentration, but were also evident at AO concentrations as low as 12.5 nM within 24 hours of transfection (**Figure 1b**). Of note, the untreated cells show diffuse nucleoplasmic NONO signal, as well as naturally occurring punctate NONO inside endogenous paraspeckle nuclear bodies (Fox *et al.* 2018) (eg. **Figure 1b(ii)**). Nuclear inclusions were also observed following 2′ O-methyl phosphorothioate AO transfection into normal human myoblasts, transformed mouse *H2K* myoblasts, and the neuroblastoma SH-SY5Y cell line (Supplementary File 1, **Figure 1)**. Once formed, the inclusions appear to be highly stable, and remain evident on the culture substrate following cell death and nuclear fragmentation and blebbing **(Figure 1c (i-ii))**. The nuclear inclusions appear as either long ‘filaments’, punctate foci, or both (**Figure 1c (iii)**). In the past we have routinely observed marked cell death associated with transfection of 2’ O-methyl phosphorothioate AOs into primary cells and cell lines, and hence explored any correlation between cell death and the formation of AO-induced nuclear inclusions. Indeed, we observed here that transfection with 2’ O-methyl phosphorothioate AO sequences induced nuclear inclusions that correlated with reduced cell survival (**Figure 1d-e**).

While transfection of AOs with phosphorothioate and phosphodiester backbones have been demonstrated to result in nuclear inclusions (Shen *et al.* 2014), PMOs are yet to be evaluated in this context. We therefore transfected fibroblasts with the AO sequences described above, synthesized as PMOs. Transfection of the PMO sequences into fibroblasts did not alter the apparent distribution of NONO, nor of any other nuclear proteins studied (**Figure 1f**). In order to demonstrate that poor cellular uptake of these compounds was not a factor, PMOs were transfected using three established delivery techniques known to lead to altered target gene expression; uncomplexed PMO (10 μM), nucleofection (1 μM), and annealed to a complementary DNA “leash” and transfected as a lipoplex (200 nM), all incubated for 72 hours after transfection (**Figure 1f (i-iii)**, respectively). Immunofluorescent detection of NONO showed no PMO-induced intranuclear inclusion formation or altered distribution of NONO following transfection, under any of the conditions used, and was reproducible, irrespective of the PMO sequence. Transfection efficiency and altered exon selection in the target *SMN2* transcript was assessed by RT-PCR across *SMN* exons 4-8, confirming that the absence of intranuclear inclusions in PMO transfected cells was not a consequence of poor uptake of PMO (**Figure 1g**).

To further explore whether AO-induced nuclear inclusions are sequence independent, ninety 2’ O-methyl phosphorothioate AOs targeting structural gene transcripts, transcription factors, splicing factors and enzymes, were transfected into fibroblasts and the cells were immunostained for SFPQ (Supplementary File 1, **Table 3)**. All AOs tested disrupted SFPQ distribution and all but one AO formed nuclear inclusions by 24 hours following AO transfection at 100 nM, irrespective of length (18-30 bases) and nucleotide composition (Supplementary File 1, **Table 3**), indicating that with optimal transfection efficiency, any 2′ O-methyl phosphorothioate sequence is likely to induce nuclear inclusions. One transfected AO induced NONO localization to the nuclear envelope (Supplementary File 1, **Figure 2a**), while a number of AO sequences induced cytoplasmic SFPQ aggregates in addition to nuclear inclusions (Supplementary File 1, **Figure 2b**). The AO-mediated cytoplasmic aggregate formation was also reported by Liang *et al.*, 2014, who observed cytoplasmic structures as a consequence of TCP1-beta subunit interaction with phosphorothioate-AOs in the cytoplasm, in addition to formation of nuclear ‘phosphorothioate bodies’, and concluded that upon transfection, the TCP1 proteins interact with phosphorothioate AOs and enhance antisense activity (in this case, activation of the RNase H-mediated target degradation) (Liang *et al.* 2015).

**Figure 2.**
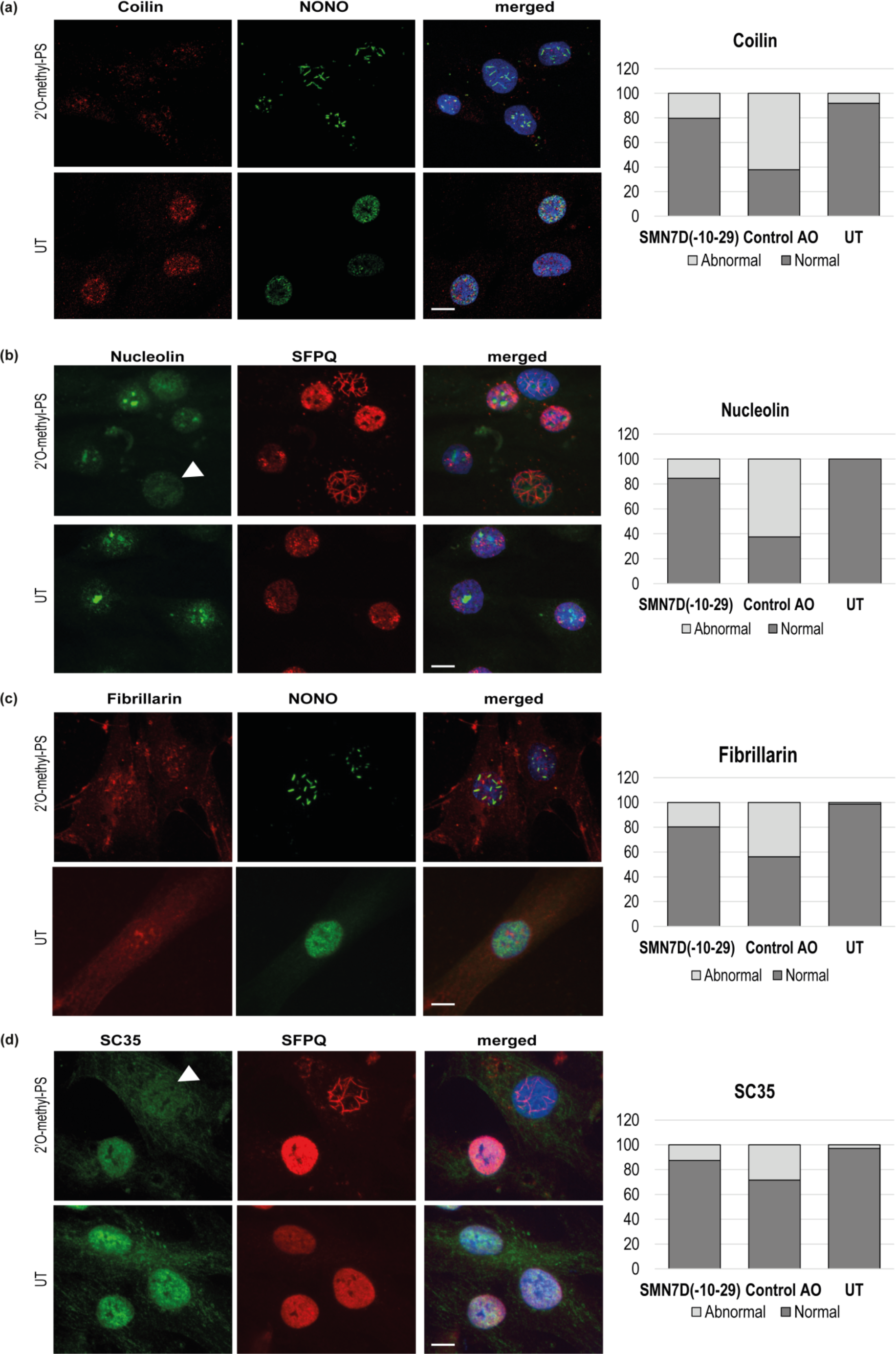
Staining of nuclear bodies following 2’ O-methyl phosphorothioate transfection (100 nM, 24 hours), with AOs as indicated in fibroblasts. **(a)** transfected with the control AO showing coilin (red) and NONO (green) overlayed with hoechst (blue), and graph showing the percentage of abnormal and normally stained cells in cultures transfected with *SMN7D(−10-29)* and control AOs; **(b)** transfected with *SMN7D(−10-29)* showing nucleolin (green) and SFPQ (red) overlayed with hoechst (blue), and graph showing the percentage of abnormal and normally stained cells in cultures transfected with *SMN7D(−10-29)* and control AOs; **(c)** transfected with the control AO showing fibrillarin (red) and NONO (green) overlayed with hoechst (blue), and graph showing the percentage of abnormal and normally stained cells in cultures transfected with with *SMN7D(−10-29)* and control AOs and **(d)** transfected with *SMN7D(−10-29)* showing SC35 (green) and SFPQ (red) overlayed with hoechst (blue), and graph showing the percentage of abnormal and normally stained cell in cultures transfected with *SMN7D(−10-29)* and control AOs. A minimum of 100 cells were counted for each sample. All scale bars = 10

### Composition of nuclear inclusions induced by 2’ O-methyl phosphorothioate AO transfection

The structure and composition of nuclear inclusions was further investigated by immunostaining of additional selected nuclear proteins found in paraspeckles, the nuclear envelope, nucleoli, nuclear speckles, Cajal bodies and nuclear stress bodies. The paraspeckle proteins PSPC1 and FUS were evident in nuclear inclusions (Supplementary File 1, **Figure 3a and 3b**) while the proteins TDP-43 and hnRNP-A1 were not observed to be components of nuclear inclusions (Supplementary File 1, **Figure 3c and 3d)**.

**Figure 3.**
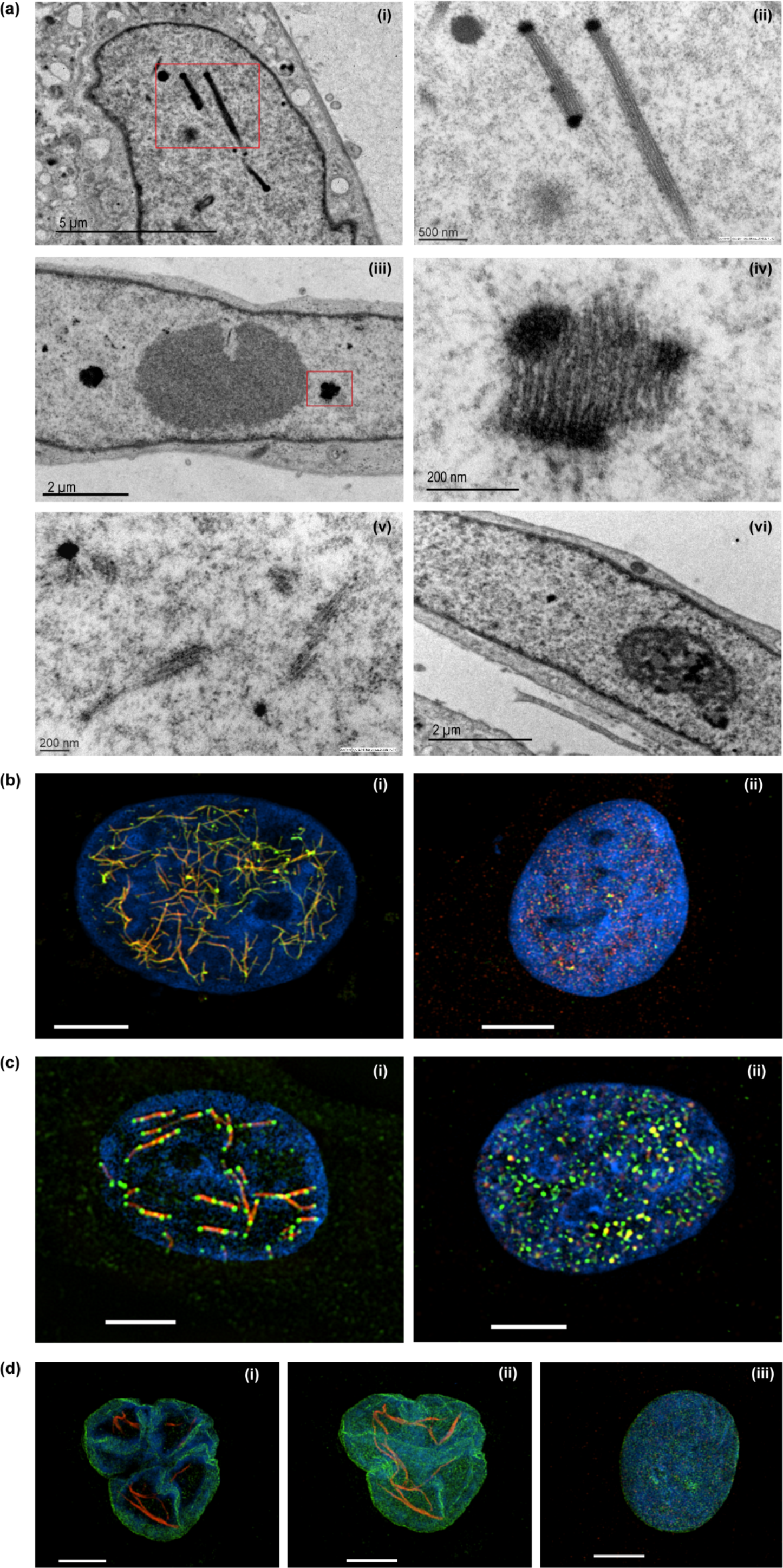
Structural analysis of nuclear inclusions in 2’ O-methyl phosphorothioate transfected fibroblasts. Fibroblasts were transfected with the 2’ O-methyl phosphorothioate AO, *SMN7D(−10-29)* at 100 nM, for 24 hours). Transfected and untreated control cells were processed for transmission electron microscopy (**a**) and super resolution fluorescence microscopy (**b-d**). Showing **(a)** transmission electron microscopy of (i) nucleus with filament-like nuclear inclusions (scale = 5 μm), (ii) higher magnification of (i) (scale = 500 nm), (iii) nucleus with foci-like nuclear inclusions (scale = 2 μm), (iv) higher magnification of (iv) (scale = 200 nm), (v) partially formed nuclear inclusions (scale = 200 nm), and (vi) untreated nucleus (scale = 2 μm); **(b)** super resolution fluorescence microscopy of SFPQ (green) and NONO (red) co-staining following 2’ O-methyl phosphorothioate (*SMN7D(−10-29))* transfection (i) and an untreated cell (ii); **(c)** super resolution fluorescence microscopy of SFPQ (red) and FUS (green) co-staining following transfection (i) and an untreated cell (ii); **(d)** super resolution fluorescence microscopy of NONO (red) and LAMIN-B1 (green) co-staining following transfection (i-ii) and an untreated cell (iii). A single z-frame is shown in (i) and a maximum intensity z-stack in (ii). For all SIM images scale bar = 5 μm.

Coilin, a component of nuclear Cajal bodies, shows altered distribution in 2′ O-methyl phosphorothioate control AO transfected cells and is distributed evenly through the nucleoplasm and cytoplasm of transfected cells showing NONO-positive nuclear inclusions, where as it is present almost exclusively in the nucleus of untreated cells (**Figure 2a)**. Nucleolin and fibrillarin, both markers for the nucleolus, show altered distribution in nuclear inclusion-containing cells, changing from nucleolar to diffuse nucleoplasmic, as shown in **Figures 2b** and **2c**, respectively. The splicing factor and component of nuclear speckles, SC35, showed enhanced cytoplasmic localisation in cells that also contained nuclear inclusions, revealed here by SFPQ immunofluorescence (**Figure 2d**).

The percentage of AO transfected cells showing disrupted staining or abnormal localization of coilin, nucleolin, fibrillarin and SC35 staining correlated with cells containing intranuclear inclusions, as indicated in the graphs (**Figure 2a-d**). Line intensity profiling shows altered distribution of these nuclear proteins in individual cells that have nuclear inclusions, and is presented in Supplementary File 1, **Figure 4**. Since components of subnuclear organelles were disorganised following transfection with phosphorothioate AOs, the levels of these proteins in the transfected cells were investigated. Western blot analysis showed that the overall abundance of each protein studied to be essentially unchanged following AO transfection. Representative western blots probed for SFPQ, NONO, TDP43, hnRNP-A1, NCL, HSF1 and beta actin, are shown in Supplementary File 1, **Figure 5**.

**Figure 4.**
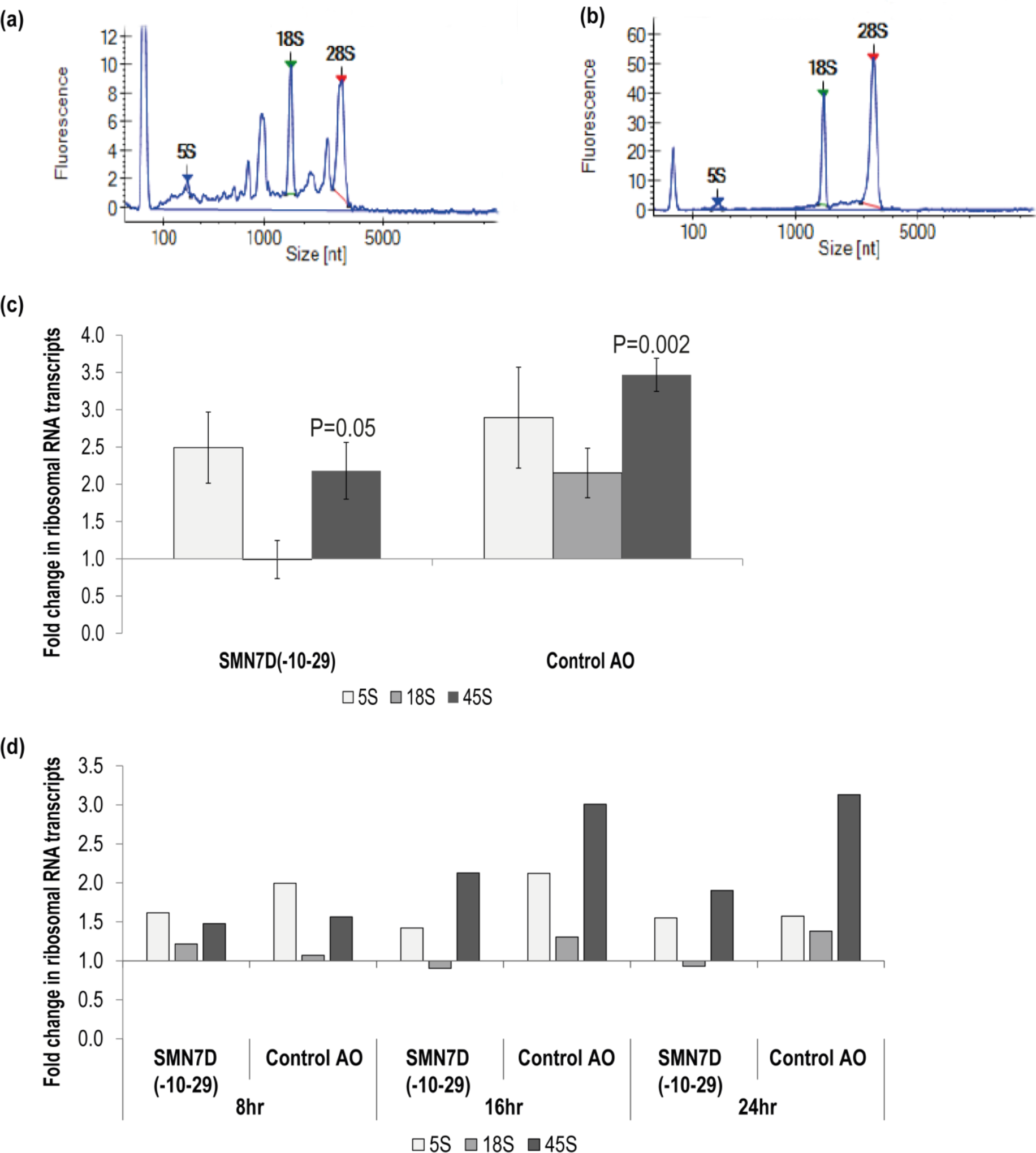
Analysis of ribosomal RNA processing. Showing bioanalyser trace of rRNA from **(a)**2’ O-methyl phosphorothioate control AO transfected cells, and **(b)** untreated cells; **(c)** qPCR analysis of 5S, 18S and 45S rRNA levels following 2’ O-methyl phosphorothioate transfection (24 hours). rRNA levels were normalised against *TBP* and compared to those in untreated cells where untreated = 1 (n=3). Error bars represent the standard error of the mean, and P-values were calculated comparing each AO treatment group to the untreated group using an unpaired T-test; and **(d)** qPCR analysis of rRNA levels over 8, 16 and 24 hours; analysis as in **(c)**(n=1).

**Figure 5.**
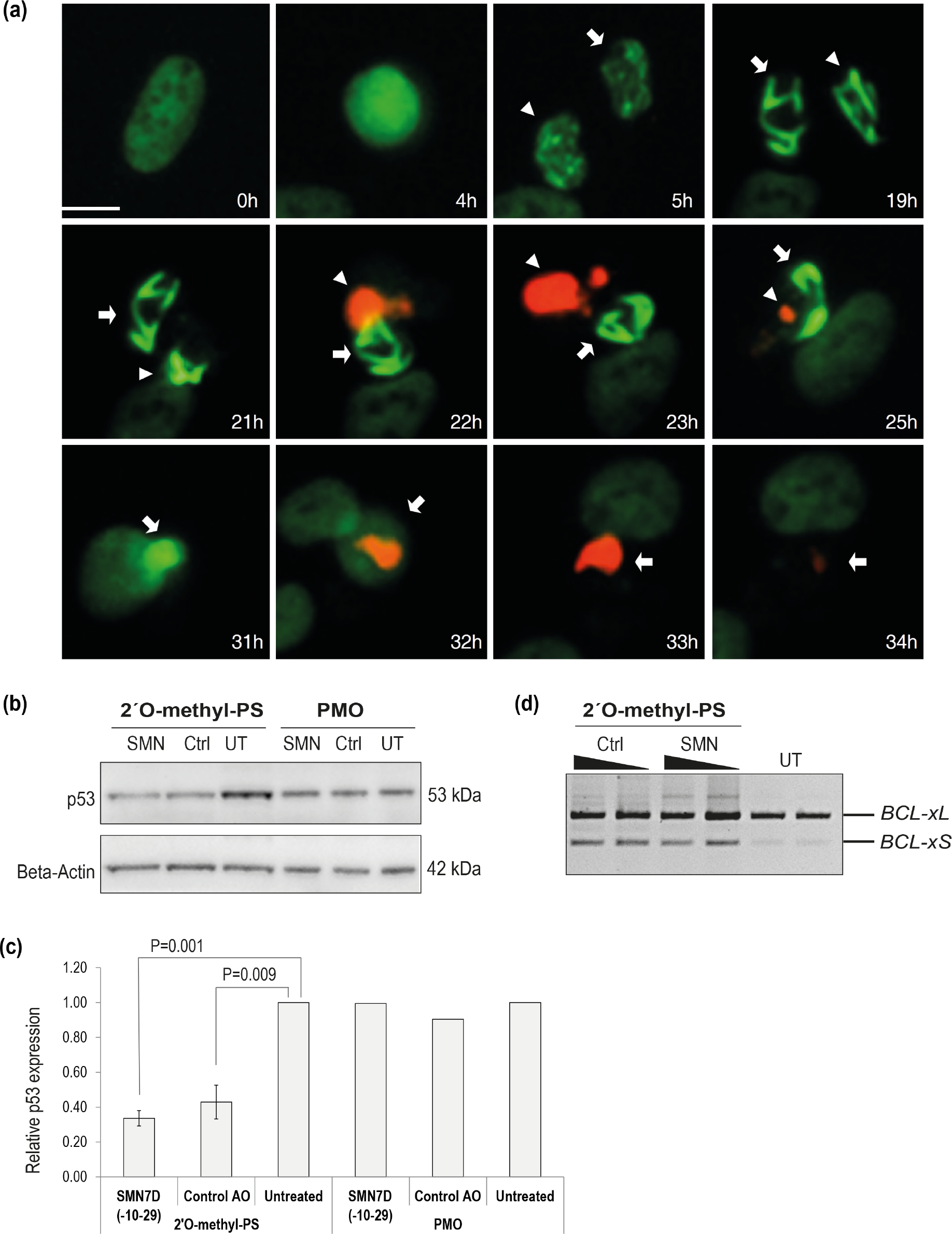
Analysis of cellular toxicity following 2’ O-methyl phosphorothioate AO transfection. **(a)** live cell imaging over time of U2OS cells with endogenous SFPQ-GFP (green) transfected with the 2’ O-methyl phosphorothioate control AO and stained for caspase 3/7 activation (red). Two sister cells are annotated with arrows, and scale bar = 10 μm; **(b)** western blot of P53 and β-actin levels in fibroblasts following 2’ O-methyl phosphorothioate AO (n=3) and PMO transfection (n=1); **(c)** densitometry analysis of **(b)** where P53 levels are normalized to β-actin and compared to those in untreated cells where untreated = 1. Error bars represent the standard error of the mean and P-values were calculated using an unpaired T-test; and **(d)** RT-PCR analysis of *BCL2* transcripts in fibroblasts following 2’O-methyl phosphorothioate AO transfection, as indicated at 100 and 50 nM (24 hours).

### Morphology of 2’ O-methyl phosphorothioate AO-induced nuclear inclusions

Detailed analysis of the 2′ O-methyl phosphorothioate *SMN7D(−10-29)*AO-induced nuclear inclusions was performed using transmission electron microscopy (TEM) on ultra-thin, osmium stained sections (**Figure 3a(i-vi))**. The nuclear inclusions appeared to be microfibrillar or amyloid-like, approximately 200-250 nm in diameter, occurring mostly in groups or bundles. For reference, the width of a DNA double helix is ~2 nm. If captured in a suitable plane of section, the structures appear to have very electron dense termini. Such nuclear inclusions have never before been revealed by transmission electron microscopy in the investigating laboratory. The electron dense regions are reminiscent of perichromatin in size and electron density, the inclusions revealed by TEM extend to ~ 2000 nm in length. Numerous small, partially formed inclusions are also evident in nuclei of transfected cells when viewed at higher magnification (**Figure 3a (iii)**).

Since transmission electron microscopy can only capture segments of the presumed large fibril-like nuclear inclusions, super resolution microscopy was used to examine 2’ O-methyl phosphorothioate *SMN7D(−10-29)* AO transfected and untreated fibroblasts (**Figure 3b-c**). AO transfected fibroblasts co-stained for NONO and SFPQ show a network of many nuclear inclusions characterised by co-localisation of the two proteins, whereas the untreated cells show more even, punctate distribution within paraspeckles (**Figure 3b (i, ii)**, respectively). Transfected fibroblasts co-stained for SFPQ and FUS reveal large interconnected fibril-like structures decorated with FUS at mid-points and at the termini (**Figure 3c (i)**). SFPQ, NONO and FUS are all sequestered to the nuclear inclusions, unlike the even nuclear distribution of these proteins in untreated cells (**Figure 3c (ii)**).

The nuclear envelope protein, lamin B1 showed altered distribution in a proportion of 2’ O-methyl phosphorothioate *SMN7D(−10-29)* transfected cells that exhibited nuclear inclusions (**Figure 3d (i-ii)**). In these cells, the nuclear envelope labelled by immunostaining of lamin B1 appears multi lobular and distorted, reminiscent of those found in the premature ageing disease progeria and in E145K cells (Taimen *et al.* 2009). A single z-frame image is shown in **Figure 3d (i)** that clearly illustrates lamin B1 localized in 4 distinct nuclear lobules. **Figure 3d** (ii and iii)shows the composite image and an image of an immunostained, untreated fibroblast. In all images (**Figure 3b-d**), the nuclear inclusions appear to reside between areas that may reflect chromosomal territories.

### 2’ O-methyl phosphorothioate AOs affect ribosomal RNA processing and maturation

Since Cajal bodies and fibrillarin are important in ribosomal RNA processing and assembly, we assessed rRNA levels in 2’ O-methyl phosphorothioate AO transfected cells and untreated cells. qPCR and bioanalyser data show unprocessed rRNA was markedly increased by 2’ O-methyl phosphorothioate AO transfection in fibroblasts (**Figure 4**). A Bioanalyser RNA trace from 2’ O-methyl phosphorothioate control AO transfected fibroblasts (**Figure 4a**) showed low levels of 18S and 28S ribosomal subunits, as well as a number of intermediate peaks compared to that on RNA from untreated fibroblasts (**Figure 4b**), consistent with incomplete rRNA processing. Poor RNA quality was ruled out, given a similar pattern was observed in all experimental repeats (n=6). This observation was supported by qPCR analysis of rRNA levels showing a 2 to 3.5-fold increase in the level of the unprocessed 45S rRNA in 2’ O-methyl phosphorothioate transfected compared to untreated fibroblasts (**Figure 4c**). Unprocessed rRNA accumulated over time, following the formation of nuclear inclusions (**Figure 4d**).

### 2’ O-methyl phosphorothioate AO transfection promotes apoptosis

Following phosphorothioate AO transfection of U2OS cells at 100 nM, live cell imaging was undertaken to evaluate caspase activation (**Figure 5a**). Nuclear inclusions were observed after 5 hours in two sister cells. Nuclear inclusions became larger and more compact overtime, followed by caspase activation after 22 hours. No caspase activation was observed in the absence of intranuclear inclusions, identifying nuclear inclusion as complicit in caspase activated apoptosis, leading to cell death. The tumour suppressor protein p53 was decreased, following transfection with 2’ O-methyl phosphorothioate AOs, relative to untreated control cells (**Figure 5b** **and c**), suggesting that the cell death observed was p53 independent, contrary to the findings by Shen *et al.*, 2018 (Shen *et al.* 2018) when using a 2′ fluoro-modified AO on a phosphorothioate backbone in the mouse. In comparison to phosphorothioate AO transfection, p53 levels remained unchanged following PMO transfection (**Figure 5b** **and c**).

The mitochondrial membrane and apoptosis regulating, *BCL2*-like gene encodes two alternatively spliced, functional protein isoforms, a longer anti-apoptotic isoform BCL-xL, and a shorter pro-apoptotic isoform BCL-xS. Increased abundance of the BCL-xS isoform induces cytochrome C release from the mitochondria, initiating caspase activated apoptosis. RT-PCR across the *BCL* transcripts (**Figure 5d**) shows 2’ O-methyl phosphorothioate AO transfection induces non-specific alternative splicing, increasing the levels of the pro-apoptotic *BCL-xS* transcript, potentially initiating the caspase activation shown in **Figure 5a**.

### Transcriptome analysis of 2’ O-methyl phosphorothioate transfected cells reveals global cellular disruptions

We next evaluated the effect of 2’ O-methyl phosphorothioate transfection and nuclear inclusion formation on the transcriptome by RNA-seq. We prepared RNA from fibroblasts transfected with the two 2’ O-methyl phosphorothioate AOs used throughout this study as well as two control samples (untreated cells and cells transfected with a DNA phosphodiester AO (DNA control) version of the same sequence as the control AO, 100 nM for 24 hours) for RNAseq analysis (n=3). The RNAseq data is available at https://www.ncbi.nlm.nih.gov/geo/query/acc.cgi?acc=GSE121713 (accession number: GSE121713). An RNAseq heatmap comparing the expression of transcripts in 2’ O-methyl phosphorothioate AO transfected fibroblasts to that in cells transfected with the DNA phosphodiester oligonucleotide and untreated cells showed stark differences in expression between the groups (**Figure 6a**). Transcripts that are overexpressed in 2’ O-methyl phosphorothioate AO transfected cells compared to controls are indicated in red, while those that are underexpressed are indicated in blue.

**Figure 6.**
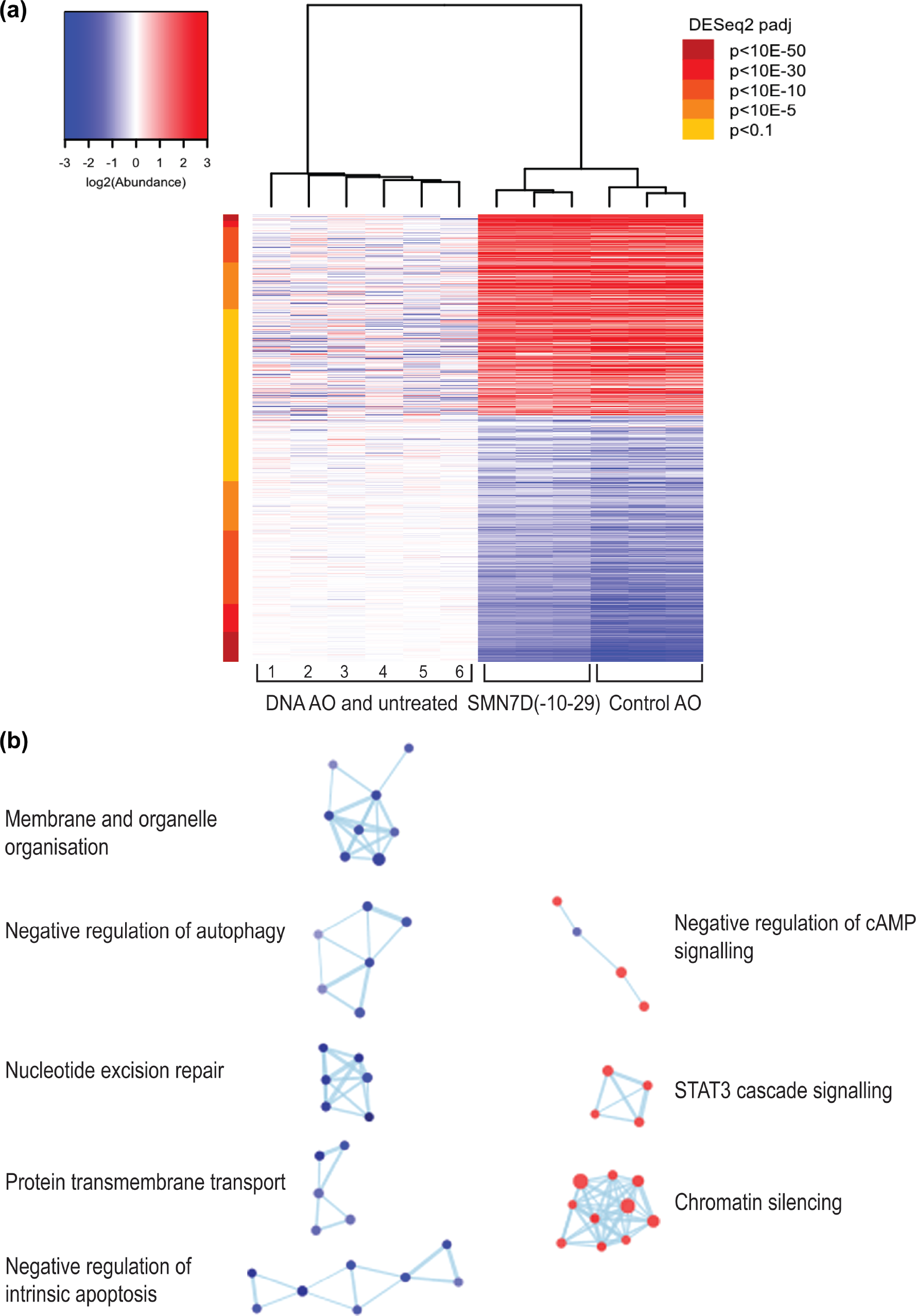
RNA sequencing analysis of 2’ O-methyl phosphorothioate AO transfected fibroblasts. **(a)** heatmap of all differentially expressed transcripts in control (1, 4, 5-untreated and 2, 3, 6, DNA AO) and 2’ O-methyl phosphorothioate AO *SMN7D(−10-29)* and control AO) transfected (100 nM, 24 hours) cells, whereby red represents overexpressed transcripts, and blue represents under-expressed transcripts; and **(b)** gene ontology (GO) network of significantly affected cellular pathways in samples from transfected, compared to untreated and DNA AO transfected cells. GO terms are represented as dots being either overexpressed (red) or under-expressed (blue), and groups with greater than 50% of genes in common are linked.

Gene ontology (GO) analyses showed significant disruptions in transcripts involved in a number of major cellular pathways. The GO network (**Figure 6b**) revealed cellular processes and pathways that are significantly over-expressed include signaling pathways, chromatin silencing, and metabolic pathways, while those that are significantly under-expressed include negative regulation of apoptosis and autophagy, membrane and organelle organisation, protein transmembrane transport and nucleotide excision repair. Gene ontology pathways included in each cellular process are listed in **Supplementary File 2**. Thus, overall these data show dramatic and widespread changes to gene expression in human fibroblasts as a result of the uptake of the 2’ O-methyl phosphorothioate AOs.

## Discussion

AOs have therapeutic potential as modulators of gene expression in many different diseases, and can do so through several different mechanisms. Antisense drugs that are in current clinical use alter exon selection during pre-mRNA processing, or induce RNaseH degradation of the target mRNA, the mechanism of action determined by the nature of the AO chemistry (for review see (Tri Le *et al.* 2016)). However, emerging reports of off-target effects conferred by synthetic oligonucleotides, identified *in vitro* (Shen *et al.* 2014, Shen *et al.* 2015) and *in vivo* (Shen *et al.* 2014, Shen *et al.* 2018), demand careful examination of the molecular effects of these compounds. Here we report that transfection of cultured human dermal fibroblasts, U2OS osteosarcoma cells, SH-SY5Y neuroblastoma cells and primary myogenic cells with 2’ O-methyl phosphorothioate oligonucleotides causes the formation of large structured nuclear inclusions, decorated with proteins that are normally components of nuclear organelles, in particular, paraspeckles. We show evidence that 2’ O-methyl phosphorothioate AO transfection alters the distribution of proteins normally associated with subnuclear bodies, impacts on global transcript expression and ribosomal RNA processing, among other critical cellular processes, and induces cell stress, followed by apoptosis.

Interaction of phosphorothioate oligonucleotides with nuclear proteins, including certain paraspeckle proteins, and the formation of paraspeckle-like structures, independent of *NEAT1*, has been reported (Shen *et al.* 2014, Shen *et al.* 2015, Shen *et al.* 2018). While Shen *et al.* (Shen *et al.* 2014) focused on the effect of phosphorothioate AOs on NONO and reported altered paraspeckle structure, this was in the context of AO-mediated modulation of gene expression, not the broader cell biology. Other off-target effects and non-specific binding of proteins, initiated by the negatively charged phosphorothioate backbone have been reported elsewhere (Dias and Stein 2002, Winkler *et al.* 2010, Liang *et al.* 2015). Hence, our experience and extensive research usage of AOs prompted further investigation of the global consequences of 2’ O-methyl phosphorothioate AO transfection on cell biology, showing many significant changes, as described above. In contrast, the transfection of (charge-neutral) PMOs in the current study did not disrupt subnuclear structures, or result in any alteration in the distribution or staining intensity of the nuclear proteins studied. This result was reproducible using multiple delivery techniques, including high dose, uncomplexed PMO transfection, transfection of PMO annealed to a DNA leash and complexed with liposome reagent, as well as nucleofection. In comparison, all 2’ O-methyl phosphorothioate AO sequences tested induced novel, structured intranuclear inclusions, revealed by transmission electron microscopy to have amyloid aggregate-like appearance.

The nuclear inclusions observed in phosphorothioate transfected cells are associated with altered nuclear architecture, disturbed gene expression and ultimately, apoptosis. While formation of such large, and apparently irreversible structures would be expected to impact on nuclear biology, the likely consequences of aberrant sequestration of paraspeckle and other nuclear proteins on RNA processing and post transcriptional regulation are of prime concern. Paraspeckles are dynamic, RNA-protein nuclear organelles that occur in the interchromatin space in mammalian cells and are now known to contain over 40 different proteins associated with one architectural long non-coding RNA, *NEAT1_2* (for review see (Fox *et al.* 2018)). Observed in most cultured cells-other than various stem cells, and cells experiencing stress, paraspeckles regulate gene expression through sequestration of proteins and RNAs (Fox *et al.* 2018), but are also implicated in microRNA biogenesis (Jiang *et al.* 2017). Paraspeckle proteins can undergo rapid, but normally reversible aggregation from monomers to form amyloid-like structures, seeded by *NEAT1_2*, and likely mediated through prion-like domains that are enriched in uncharged polar amino acids (Lancaster *et al.* 2014). The mechanism by which phosphorothioate oligonucleotides induce some paraspeckle proteins to form aberrant aggregates is unknown, however, we do know that sub cellular localization and phosphorothioate AO behaviour in transfected cells is influenced by protein interactions and the 2’ sugar AO modifications (Bailey *et al.* 2017, Crooke *et al.* 2017), and that synthetic double stranded RNA is also able to seed *de novo* paraspeckle formation (Shelkovnikova *et al.* 2018). How other nuclear proteins become incorporated into the phosphorothioate AO induced nuclear inclusions, and why TDP43, a major component of paraspeckles does not appear in these inclusions in our study, remains elusive.

Immunofluorescent staining of 2’ O-methyl phosphorothioate transfected fibroblasts revealed the paraspeckle components NONO, PSPC1, SFPQ and FUS co-localised with large nuclear structures in excess of 2000 nm in length, with FUS decorating the structures in an ordered manner. All four of these proteins include prion-like domains (for review see (Fox *et al* 2018)) and we speculate that the known propensity of phosphorothioate backbone compounds to interact with and bind proteins then alters the liquid-liquid phase properties of the paraspeckle proteins, shifting them towards insoluble, amyloid-like aggregations. However, not all paraspeckle components with prion-like domains investigated were found to colocalise with the inclusions, and in addition, the localisation and distribution of some proteins associated with other subnuclear bodies was altered in cells with nuclear inclusions.

Coilin, an integral component of Cajal bodies, nucleolin, one of the most abundant proteins in the nucleolus but also found in the cytoplasm and on the cell membrane, fibrillarin, located in the dense fibrillar component of the nucleolus and SC35 (SRSF2), an essential splicing factor found in nuclear speckles all showed altered distribution in cells that have phosphorothioate AO-induced nuclear inclusions, but did not co-localize with these inclusions. Cajal bodies assemble spliceosomal and nucleolar ribonucleoproteins required for pre-mRNA and pre-rRNA processing, and are recently proposed to contribute to genome organization, with global effects on gene expression and RNA splicing (Wang *et al.* 2016). The nucleolus is the most prominent nuclear structure, and is where synthesis and processing of ribosomal transcripts to yield the mature rRNAs *5.8S, 18S* and *28S* from the *45S* pre-rRNA takes place. We show that ribosomal RNA processing is greatly impaired by 2’ O-methyl phosphorothioate AO transfection, and speculate that this is a likely manifestation of improper localization of major protein components of the nucleolus and Cajal bodies and consequent disruption to their functions. In addition, the aberrant inclusions that we speculate occupy interchromosomal spaces may well impose physical constraints upon nuclear organisation, and prevent proper localisation of Cajal bodies and the nucleoli close to their normal chromosomal sites. Tissues with high demand for transcript splicing and ribosome biogenesis, and neurons in particular, have prominent Cajal bodies, juxtaposed to nucleoli (for review see (Lafarga *et al.* 2017)), and disruption or loss of Cajal bodies is associated with severe neuronal dysfunction (Tapia *et al.* 2017). Indeed, disruption or depletion of Cajal bodies was seen as the earliest nuclear sign of motor neuron degeneration in a spinal muscular atrophy mouse model and induced a progressive nucleolar dysfunction in ribosome biogenesis (Tapia *et al.* 2017).

Paraspeckle biology has gained increasing interest due to the association of paraspeckle proteins with neurodegenerative disease, in particular amyotrophic lateral sclerosis (ALS), and the observation that *NEAT1_2* is up-regulated in early-stage motor neurons from the spinal cords of ALS patients (Shelkovnikova *et al.* 2014, Yamazaki and Hirose 2015). Paraspeckle formation, not observed in healthy spinal motor neurons, is enhanced in spinal cords of patients with early stage sporadic and familial ALS, and mutations in many paraspeckle proteins (eg. TDP-43, FUS, NONO, SFPQ) are associated with ALS (Nishimoto *et al.* 2013, Shelkovnikova *et al.* 2018). Whether paraspeckles are protective or causative in ALS molecular pathology is not known at this time (Yamazaki and Hirose 2015), nor is the role of paraspeckles in disease well understood. Nevertheless, phosphorothioate AO-mediated dysregulation of paraspeckle formation and altered nucleoli and Cajal body biology shown in our study, together with the building body of evidence that nuclear organelle dysfunction (Lafarga *et al.* 2017, Tapia *et al.* 2017) has implications for central nervous system, and probably all, clinical applications of these compounds.

Not surprisingly, considering the altered nuclear architecture and distribution of proteins implicated in RNA biology, transcriptome sequencing showed significant global effects on the expression of transcripts and revealed disturbances to many critical cellular processes in phosphorothioate AO transfected cells. Of particular concern, pathways involved in apoptosis, chromatin silencing, cellular metabolism and a number of signalling pathways, autophagy and nucleotide excision repair were disturbed. While we acknowledge that these *in vitro* studies in replicating cells may not fully reflect potential off-target treatment effects in tissues *in vivo*, Toonan et al., 2018 (Toonen *et al.* 2018) report significant upregulation of immune system-associated genes in brains of mice treated by intracerebroventricular injection of 2’ O-methyl phosphorothioate AO. The upregulation of immune system associated genes was detectable for at least 2 months after the last AO administration. Here, the exaggerated sequestration of nuclear proteins as a result of 2’ O-methyl phosphorothioate AO transfection dramatically disturbs RNA processing, disrupts nuclear architecture, and induces apoptosis, and it seems reasonable to consider that injection site reactions and adverse effects reported after clinical evaluation of the 2’ O-methyl phosphorothioate *Drisapersen* for the treatment of Duchenne muscular dystrophy (Mendell *et al.* 2017) and *Kynamro^®^* (*Mipomersen*) for the treatment of familial hypercholesterolemia (Wong and Goldberg 2014) may be mediated at least in part, by non-specific interactions of nuclear components with the AO backbone. It might also be prudent to deliberate on the reported effects of antisense drugs, attributed to the AO action on the target transcript, and consider whether some level of the apparent antisense effect on splicing, in particular, could perhaps be a consequence of disturbance of RNA processing pathways more broadly.

We speculate that, unlike endogenous paraspeckles and other dynamic nuclear bodies, the formation of the aberrant nuclear inclusions seen *in vitro* here does not appear to be reversible. While 2’ O-methyl phosphorothioate transfected cultures showed reduced cell numbers, whether this was due to cell death or impaired replication, or both, is uncertain, and whether the nuclear inclusions *per se* or reduced availability of RNA processing are primarily responsible requires further investigation. However, any effects resulting in perturbation of nuclear proteins may be compounded by the formation of the nuclear amyloid-like aggregates and likely disturbance of protein homeostasis, termed ‘proteostasis’ (Yerbury *et al.* 2016). Although other descriptions of exogenously induced nuclear amyloid-like aggregates are limited, Arnhold *et al.*, 2015 (Arnhold *et al.* 2015) identified large amyloid-like aggregates in SH-SY5Y cells, treated with mercury, a notorious neurotoxicant, in a study that explains the mechanism of heavy metal neurotoxicity and identified amyloid protein aggregation in the cell nucleus as causative. Mass spectrometry of the purified protein aggregates identified a subset of spliceosomal components and the nuclear envelope protein lamin B1 (Arnhold *et al.* 2015). In our study, we also detected changes in lamin B1, although here we did not detect lamin B1 in the nuclear inclusions, we nevertheless observed that nuclear membrane and lamin B1 organization was distorted in fibroblasts containing nuclear inclusions. Interestingly, the multi-lobulated nuclei and lamin B1 staining are reminiscent of cells from progeria patients carrying the lamin A 433G>A mutation (E145K) (Taimen *et al.* 2009).

In summary, we report phosphorothioate backbone-specific effects of modified oligonucleotides on the distribution and localization of nuclear proteins, appearance of novel nuclear structures composed in part of a subset of paraspeckle protein components, and sequence-independent effects on nascent RNA processing. While the *in vivo*, longer term repercussions of exogenous oligonucleotide-induced nuclear protein aggregates that include many paraspeckle components and cause sub-nuclear disorganisation are yet to be determined, our observations suggest that phosphorothioate backbone antisense compounds destined for clinical application would benefit from further scrutiny.

## Acknowledgments

The authors would like to acknowledge technical advice from Russell Johnsen. This work was performed in part during LLF’s and ILP’s PhD candidatures. Laboratory studies and ILP were supported by funding from the Parry Foundation through the Spinal Muscular Atrophy Association of Australia and LLF received a Team Spencer and Muscular Dystrophy WA PhD stipend. The authors received research support from the NHMRC Project Grants APP1147496, APP1086311, APP1144791 and the MNDi foundation, Western Australia.

## Author Contributions

Conceived and designed experiments: LLF, RL, SF, MTA, ILP and AF. Performed experiments: LLF, RL, MTA, ILP, AH, and LG. Analysis of RNA-seq data: JC. Prepared and edited manuscript: LLF, SF, RL, MTA, ILP, JC, AH, LG, SDW and AF.

## Abbreviations

AO: antisense oligonucleotide
SMA: spinal muscular atrophy
DMD: Duchenne muscular dystrophy
PMO: phosphorodiamidate morpholino oligomer
ALS: amyotrophic lateral sclerosis

