## Supplementary File 1 for "Interaction of modified oligonucleotides with nuclear proteins, formation of novel nuclear structures and sequence-independent effects on RNA processing"

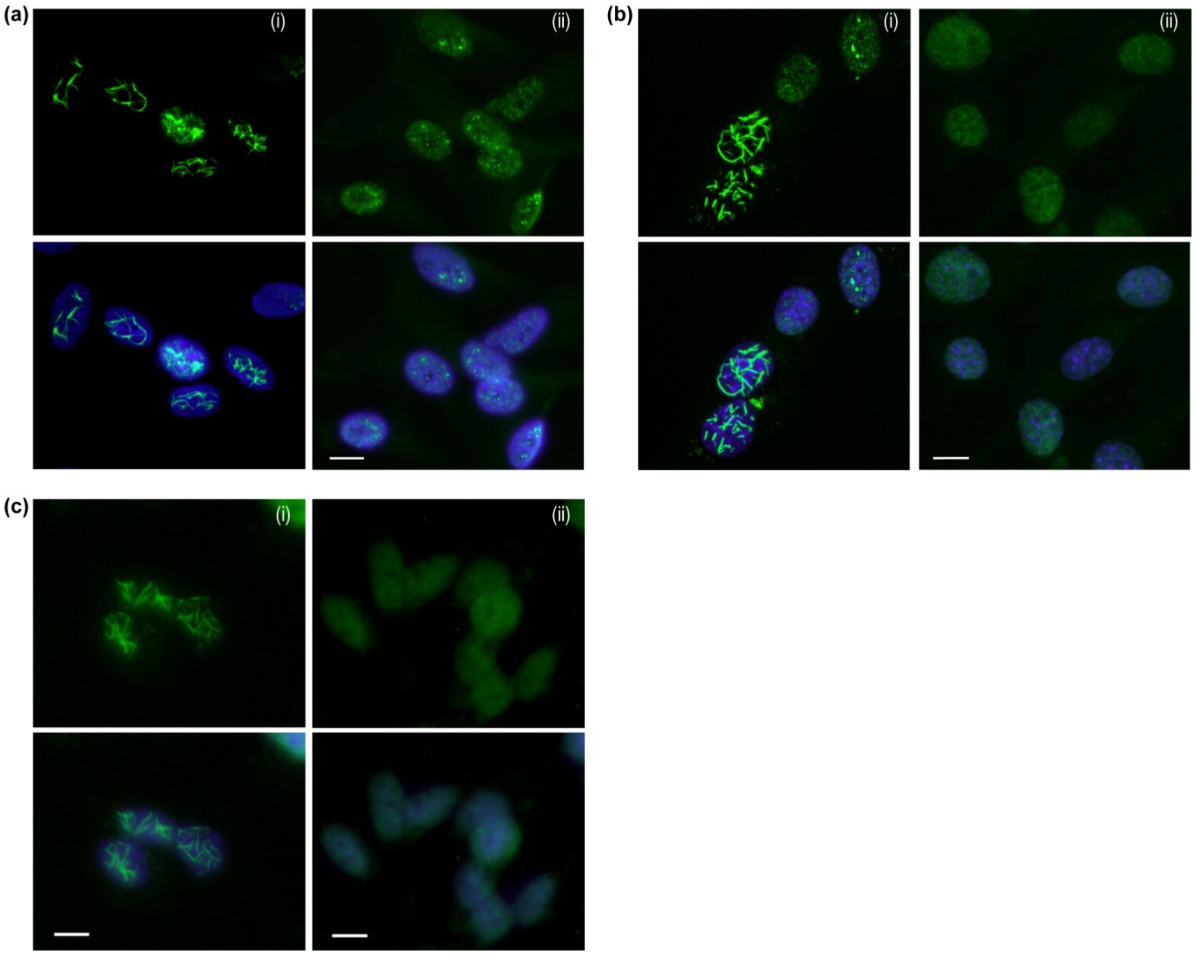

**Supplementary Figure 1 – Immunofluorescence staining of paraspeckle proteins in additional cell types**. Staining was carried out following transfection with **(a)** (i) 2´ O-methyl phosphorothioate *SMN7D(-10-29)* in primary human myogenic cells; **(b)** (i) 2´ O-methyl phosphorothioate AO targeting *Smn* in mouse myogenic cells; and **(c)** (i) the 2´ O-methyl phosphorothioate control AO in SH-SY5Y neuroblastoma cells. (i) 2´ O-methyl phosphorothioate-transfected and (ii) untreated shown in each panel. NONO was immunostained in human cells and SFPQ in mouse cells. Scale bar = 10 μm.

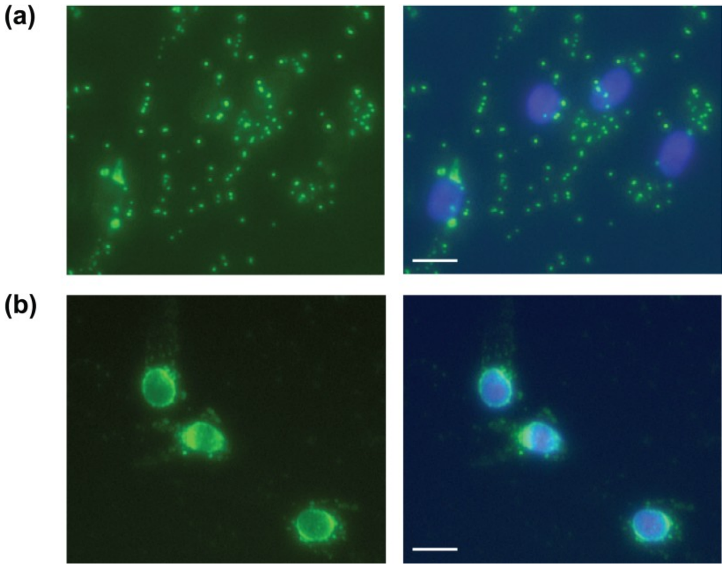

**Supplementary Figure 2 – Immunofluorescence imaging of alternative paraspeckle protein sequestration and staining patterns.** Overlay of Hoechst and SFPQ staining in fibroblasts following 2´O-methyl phosphorothioate transfection (100 nM, 24 hours), showing **(a)** cytoplasmic SFPQ staining in cells without nuclear inclusions after transfection with AO 43 (Supplementary Table 3); and **(b)** staining of SFPQ around the nuclear envelope in cells without nuclear inclusions, transfected with AO 90 (Supplementary Table 3). Scale bar = 10 µm.

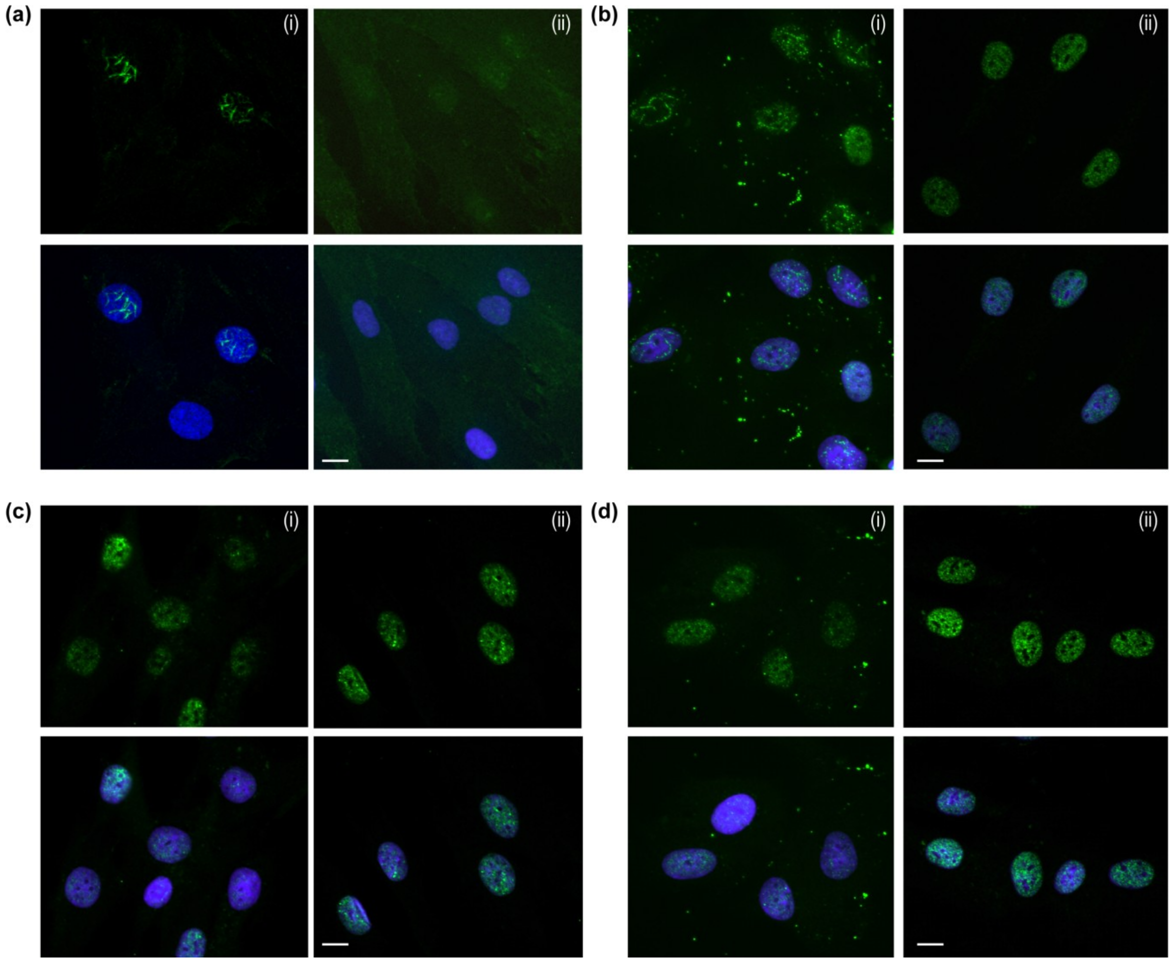

**Supplementary Figure 3 - Immunofluorescence staining of additional paraspeckle proteins following 2´ O-methyl phosphorothioate *SMN7D(-10-29)* transfection (100 nM for 24 hours)**. Showing **(a)** paraspeckle protein component 1 (PSPC1); **(b)** fused in sarcoma (FUS); **(c)** TAR-DNA binding protein 43 (TDP43); and **(d)** heterogeneous nuclear ribonucleoprotein A1 (hnRNPA1); in (i) 2´ O-methyl transfected and (ii) untreated cells shown in each panel. Scale bar = 10 μm.

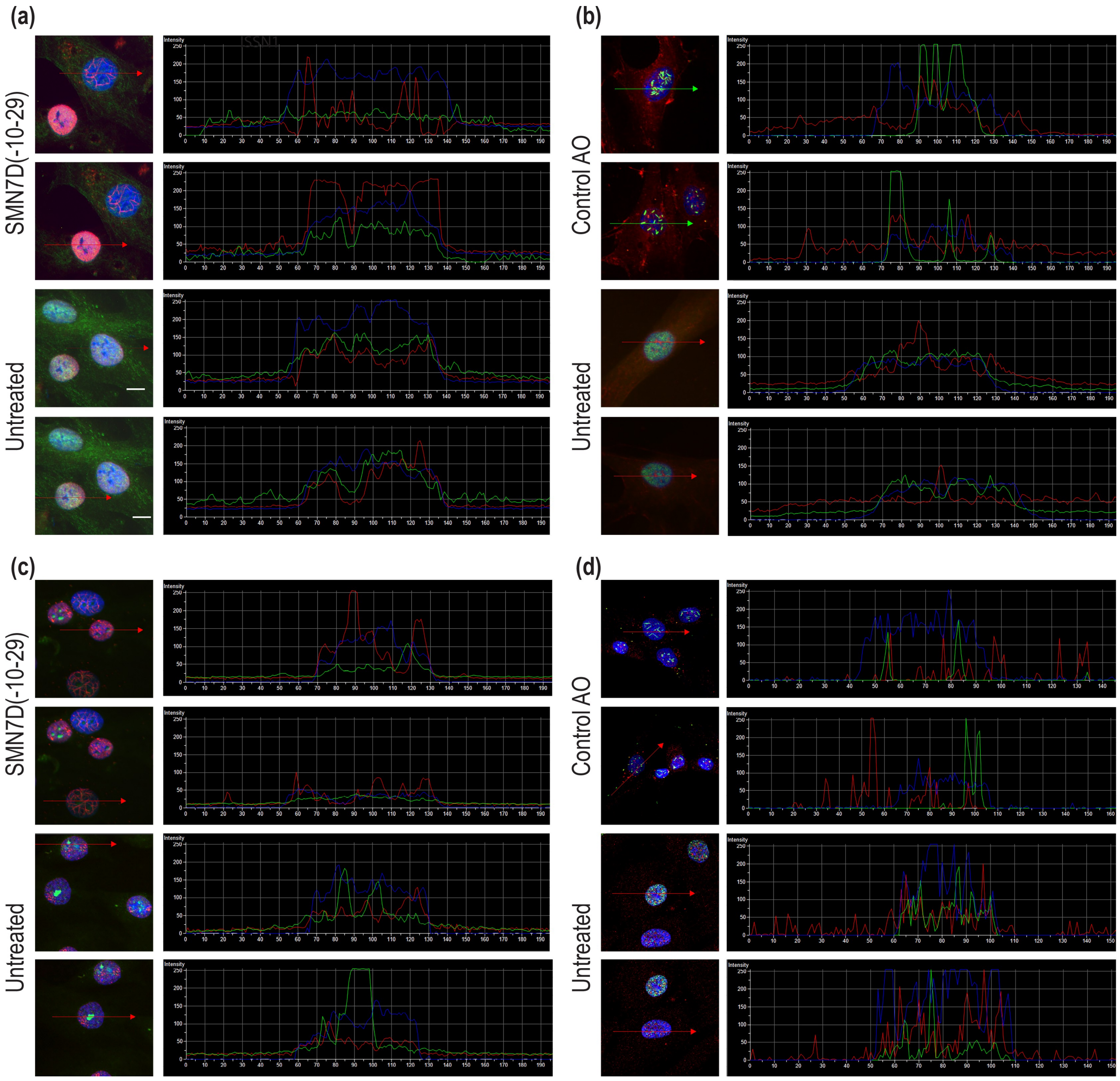

**Supplementary Figure 4. – Intensity profiling of immunofluorescent staining of paraspeckle proteins following transfection of the 2´ O- methyl phosphorothioate AOs.** Including, *SMN7D(-10-29)* **(a, c)** and the Control AO **(b, d)** (100 nM for 24 hours) and intensity profiling of selected cells showing **(a)** SFPQ (red) and SC35 (green); **(b)** NONO (green) and FBL (red); **(c)** SFPQ (red) and NCL (green); **(d)** NONO (green) and coilin (red).

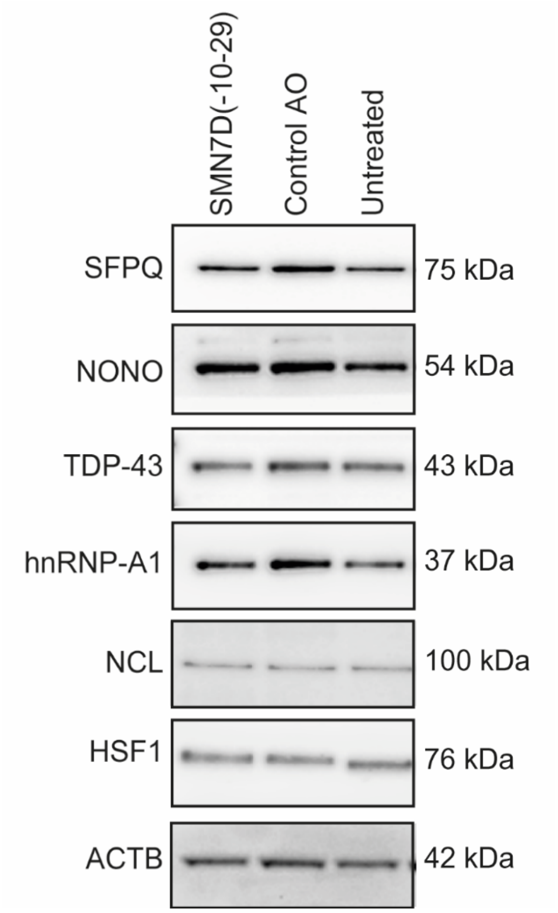

**Supplementary Figure 5 – Western blots following 2´ O-methyl phosphorothioate AO transfection.** Western blots of extracts from cells transfected with the *SMN7D(-10-29)* and Control AOs (100 nM for 24 hours), probed with antibodies that recognize SFPQ, NONO, TDP-43, hnRNP-A1, NCL, HSF1 and ACTB (Supplementary Table 2).

**Supplementary Tables:**

**Supplementary Table 1** primer sequences and PCR conditions used in this study.

| Primers | Concentration/amount of primer per reaction | Sequence 5´-3´ | Temperature profile |
|---|---|---|---|
| *SMN* | 25 ng | AGGTCTCCTGGAAATAAATCAG<br>TGGTGTCATTTAGTGCTGCTCT | 55°C 30 min<br>94°C 2 min<br>25 cycles:<br>94°C 40 sec<br>56°C 30 sec<br>68°C 1 min |
| *BCL2* | 25 ng | AAT GTC TCA GAG CAA CCG GG<br>GGG AGG GTA GAG TGG ATG GT | 55°C 30 min<br>94°C 2 min<br>25 cycles:<br>94°C 40 sec<br>60°C 30 sec<br>68°C 1 min |
| *5S* | 50 nM | GGC CAT ACC ACC CTG AAC GC<br>CAG CAC CCG GTA TTC CCA GG | 95 °C 20 sec<br>40 cycles<br>95 °C 3 sec<br>60 °C 30 sec |
| *18S* | 250 nM | GTA ACC CGT TGA ACC CCA TT<br>CCA TCC AAT CGG TAG TAG CG | |
| *45S* | 100 nM | TGT CAG GCG TTC TCG TCT C<br>AGC ACG ACG TCA CCA CAT C | |
| *TBP* | 500 nM | TCAGGCGTTCGGTGGATCGAGT<br>AGTGATGCTGGGCACTGCGGAGAA | |
| *TUBB* | 100 nM | CTT CGG CCA GAT CTT CAG AC<br>AGA GAG TGG GTC AGC TGG AA | |

**Supplementary Table 2** Antibodies used in this study, indicating dilutions used, detection method and antibody validation.

| Protein | Protein name | Supplier | Catalogue # | Dilution | | Conjugate | Validation |
|---|---|---|---|---|---|---|---|
| | | | | IF | WB | | |
| NONO | Non-POU domain containing octamer | Prepared in house | NA | 1:1000 | 1:10000 | mouse | (Souquere, Beauclair et al. 2010) |
| SFPQ | Splicing factor proline & glutamine rich | Abcam | Ab38148 | 1:1000 | 1:8000 | rabbit | (Ke, Dramiga et al. 2012) |
| PSPC1 | paraspeckle protein component 1 | Merck Millipore | HPA038904 | 1:50 | NA | mouse | Antibodypedia WB, ICC, IHC |
| FUS | Fused in sarcoma | Santa Cruz | Sc-47711 | 1:500 | NA | mouse | Validated in SW480 cells (https://datasheets.scbt.com/sc-47711.pdf) |
| TDP-43 | TAR-DNA binding protein 43 | Protein Tech | 10782-2-AP | 1:300 | 1:3000 | rabbit | Antibodypedia WB, ICC, IHC |
| hnRNP-A1 | Heterogeneous nuclear ribonucleoprotein A1 | Thermo Scientific | PA5-19431 | 1:700 | 1:2000 | rabbit | Antibodypedia WB, IHC |
| NCL | Nucleolin | Thermo Scientific | 39-6400 | 1:250 | 1:2000 | mouse | Antibodypedia WB, ICC |
| FBL | Fibrillarin | Cell Signalling | 2639 | 1:400 | NA | rabbit | Antibodypedia WB, ICC |
| COIL | Coilin | Sapphire Bioscience | GTX112570 | 1:250 | NA | rabbit | Validated in HeLa, HEPG2 and 293T cells, and in cell extracts (http://www.genetex.com/Coilin-antibody-GTX112570.html) |
| LMNB1 | Lamin B1 | Protein Tech | 12987-1-AP | 1:500 | NA | rabbit | Antibodypedia WB, ICC, IHC |

| | | | | | | | |
|---|---|---|---|---|---|---|---|
| SC35 | Splicing factor SC35 | Merk Millipore | 04-1550 | 1:1000 | NA | mouse | (Werwein, Dzuganova et al. 2013) |
| HSF1 | Heat shock factor 1 | Cell Signaling | 12972 | 1:400 | 1:500 | rabbit | Antibodypedia WB, ICC, IHC |
| ACTB | Beta actin | Sigma | A5441 | NA | 1:60000 | mouse | Validated in HS-68(WB), HeLa, JURKAT, COS7, NIH-3T3, PC-12, RAT2, CHO, MDBK, MDCK, FS-11 human fibroblast (ICC) https://www.sigmaaldrich.com/catalog/product/sigma/a5441?lang=en&region=AU |
| P53 | Tumor Protein 53 | Novocastra | NCL-Lp53-D07 | NA | 1:1000 | mouse | (Horne, Anderson et al. 1996) (WB) |

**Supplementary Table 3** Results of SFPQ immunostaining following transfection of fibroblasts with 2′ O-methyl phosphorothioate AOs (100 nM, 24 hours). The nucleotide composition and length of each AO is indicated. Cells were stained for SFPQ and the percentages of cells with nuclear inclusions recorded, as was the percentage of cells showing cytoplasmic SFPQ aggregation.

| AO # | Nucleotide composition | | | | Length | Cells with nuclear inclusions | Total cell number | SFPQ-nuclear inclusions (% cells) | Cytoplasmic SFPQ staining | Comments |
|---|---|---|---|---|---|---|---|---|---|---|
| | A | C | G | U | | | | | | |
| 1 | 4 | 9 | 4 | 8 | 25 | 56 | 56 | **100.0** | | |
| 2 | 5 | 9 | 3 | 8 | 25 | 60 | 60 | **100.0** | | |
| 3 | 3 | 8 | 3 | 11 | 25 | 107 | 109 | **98.2** | | |
| 4 | 4 | 10 | 3 | 8 | 25 | 87 | 89 | **97.8** | | |
| 5 | 4 | 8 | 7 | 6 | 25 | 118 | 121 | **97.5** | >90% | |
| 6 | 5 | 11 | 4 | 5 | 25 | 95 | 98 | **96.9** | | |
| 7 | 6 | 7 | 8 | 4 | 25 | 29 | 30 | **96.7** | | |
| 8 | 4 | 9 | 3 | 9 | 25 | 112 | 116 | **96.6** | | |
| 9 | 4 | 7 | 3 | 11 | 25 | 76 | 79 | **96.2** | | |
| 10 | 7 | 2 | 7 | 9 | 25 | 70 | 73 | **95.9** | | |
| 11 | 6 | 2 | 3 | 14 | 25 | 128 | 134 | **95.5** | | |
| 12 | 7 | 8 | 3 | 7 | 25 | 63 | 66 | **95.5** | | |
| 13 | 7 | 7 | 3 | 8 | 25 | 55 | 58 | **94.8** | | |
| 14 | 10 | 10 | 5 | 2 | 27 | 78 | 84 | **92.9** | | |
| 15 | 10 | 7 | 2 | 6 | 25 | 130 | 140 | **92.9** | | |
| 16 | 4 | 9 | 2 | 9 | 24 | 122 | 132 | **92.4** | | |
| 17 | 11 | 3 | 9 | 7 | 30 | 101 | 110 | **91.8** | | |
| 18 | 5 | 10 | 4 | 6 | 25 | 52 | 57 | **91.2** | | |
| 19 | 4 | 6 | 8 | 7 | 25 | 29 | 32 | **90.6** | | |
| 20 | 1 | 12 | 0 | 12 | 25 | 33 | 37 | **89.2** | | |
| 21 | 15 | 2 | 3 | 2 | 22 | 44 | 50 | **88.0** | | |
| 22 | 11 | 3 | 10 | 2 | 26 | 109 | 124 | **87.9** | >90% | |
| 23 | 6 | 6 | 4 | 4 | 20 | 95 | 111 | **85.6** | | |
| 24 | 7 | 8 | 2 | 8 | 25 | 53 | 63 | **84.1** | | |
| 25 | 4 | 9 | 2 | 11 | 26 | 118 | 142 | **83.1** | | |
| 26 | 7 | 8 | 3 | 7 | 25 | 90 | 109 | **82.6** | | |
| 27 | 5 | 7 | 8 | 6 | 26 | 98 | 119 | **82.4** | | |
| 28 | 3 | 9 | 8 | 5 | 25 | 98 | 120 | **81.7** | | |
| 29 | 3 | 14 | 6 | 3 | 26 | 40 | 50 | **80.0** | | |
| 30 | 3 | 7 | 7 | 8 | 25 | 57 | 72 | **79.2** | | |
| 31 | 0 | 8 | 3 | 14 | 25 | 88 | 113 | **77.9** | | |
| 32 | 6 | 9 | 5 | 5 | 25 | 109 | 140 | **77.9** | | |
| 33 | 18 | 2 | 6 | 3 | 29 | 100 | 129 | **77.5** | | |
| 34 | 6 | 7 | 5 | 7 | 25 | 62 | 81 | **76.5** | | |
| 35 | 3 | 10 | 3 | 12 | 28 | 104 | 136 | **76.5** | | |
| 36 | 14 | 5 | 5 | 4 | 28 | 26 | 35 | **74.3** | | |

| | | | | | | | | | |
|---|---|---|---|---|---|---|---|---|---|
| 37 | 3 | 9 | 6 | 7 | 25 | 56 | 77 | **72.7** | |
| 38 | 10 | 7 | 4 | 4 | 25 | 52 | 72 | **72.2** | |
| 39 | 9 | 8 | 4 | 4 | 25 | 114 | 159 | **71.7** | >90% |
| 40 | 5 | 7 | 5 | 8 | 25 | 77 | 108 | **71.3** | |
| 41 | 1 | 9 | 4 | 4 | 18 | 72 | 109 | **66.1** | |
| 42 | 8 | 4 | 9 | 4 | 25 | 83 | 126 | **65.9** | >90% |
| 43 | 3 | 10 | 8 | 4 | 25 | 86 | 131 | **65.6** | >90% |
| 44 | 9 | 3 | 10 | 3 | 25 | 75 | 115 | **65.2** | >90% |
| 45 | 8 | 4 | 7 | 6 | 25 | 47 | 74 | **63.5** | |
| 46 | 6 | 7 | 2 | 10 | 25 | 100 | 158 | **63.3** | |
| 47 | 10 | 3 | 5 | 7 | 25 | 65 | 103 | **63.1** | |
| 48 | 7 | 6 | 10 | 2 | 25 | 72 | 115 | **62.6** | >90% |
| 49 | 7 | 7 | 8 | 11 | 33 | 25 | 40 | **62.5** | |
| 50 | 6 | 4 | 8 | 2 | 20 | 83 | 137 | **60.6** | >90% |
| 51 | 4 | 8 | 6 | 7 | 25 | 54 | 93 | **58.1** | |
| 52 | 1 | 10 | 11 | 8 | 30 | 36 | 62 | **58.1** | |
| 53 | 2 | 7 | 8 | 8 | 25 | 78 | 138 | **56.5** | |
| 54 | 6 | 8 | 6 | 5 | 25 | 55 | 100 | **55.0** | |
| 55 | 7 | 5 | 8 | 4 | 24 | 68 | 125 | **54.4** | |
| 56 | 8 | 4 | 7 | 6 | 25 | 60 | 112 | **53.6** | |
| 57 | 5 | 7 | 6 | 7 | 25 | 40 | 77 | **51.9** | |
| 58 | 6 | 9 | 6 | 5 | 26 | 95 | 187 | **50.8** | |
| 59 | 8 | 4 | 5 | 8 | 25 | 34 | 67 | **50.7** | |
| 60 | 6 | 5 | 6 | 8 | 25 | 99 | 207 | **47.8** | |
| 61 | 6 | 3 | 13 | 3 | 25 | 23 | 49 | **46.9** | >90% |
| 62 | 9 | 4 | 9 | 3 | 25 | 41 | 89 | **46.1** | >90% |
| 63 | 8 | 7 | 7 | 3 | 25 | 55 | 121 | **45.5** | |
| 64 | 6 | 1 | 6 | 11 | 24 | 65 | 146 | **44.5** | |
| 65 | 3 | 7 | 8 | 7 | 25 | 57 | 134 | **42.5** | |
| 66 | 11 | 9 | 3 | 8 | 31 | 44 | 104 | **42.3** | |
| 67 | 3 | 10 | 7 | 5 | 25 | 22 | 55 | **40.0** | |
| 68 | 6 | 7 | 9 | 3 | 25 | 48 | 122 | **39.3** | >90% |
| 69 | 2 | 9 | 13 | 1 | 25 | 43 | 114 | **37.7** | >90% |
| 70 | 2 | 7 | 6 | 4 | 19 | 45 | 123 | **36.6** | |
| 71 | 6 | 7 | 9 | 3 | 25 | 36 | 104 | **34.6** | |
| 72 | 5 | 6 | 3 | 0 | 14 | 45 | 132 | **34.1** | |
| 73 | 9 | 7 | 5 | 4 | 25 | 30 | 90 | **33.3** | |
| 74 | 6 | 3 | 7 | 9 | 25 | 29 | 88 | **33.0** | >90% |
| 75 | 5 | 10 | 8 | 2 | 25 | 52 | 160 | **32.5** | |
| 76 | 5 | 6 | 4 | 5 | 20 | 17 | 59 | **28.8** | |
| 77 | 6 | 6 | 8 | 1 | 21 | 33 | 115 | **28.7** | |
| 78 | 5 | 6 | 7 | 7 | 25 | 33 | 117 | **28.2** | |
| 79 | 3 | 8 | 5 | 9 | 25 | 39 | 143 | **27.3** | |
| 80 | 3 | 5 | 12 | 5 | 25 | 26 | 96 | **27.1** | >90% |
| 81 | 7 | 8 | 7 | 3 | 25 | 21 | 78 | **26.9** | |
| 82 | 6 | 4 | 9 | 6 | 25 | 32 | 122 | **26.2** | >90% |

| | | | | | | | | | | |
|---|---|---|---|---|---|---|---|---|---|---|
| 83 | 4 | 10 | 6 | 5 | 25 | 40 | 156 | **25.6** | | |
| 84 | 4 | 8 | 10 | 2 | 24 | 30 | 131 | **22.9** | | |
| 85 | 4 | 5 | 9 | 5 | 23 | 15 | 134 | **11.2** | >90% | |
| 86 | 4 | 3 | 8 | 4 | 19 | 10 | 138 | **7.2** | >90% | |
| 87 | 2 | 4 | 11 | 8 | 25 | 8 | 146 | **5.5** | >90% | |
| 88 | 6 | 6 | 12 | 3 | 27 | 3 | 169 | **1.8** | >90% | |
| 89 | 9 | 4 | 11 | 1 | 25 | 2 | 161 | **1.2** | >90% | |
| 90 | 3 | 4 | 12 | 6 | 25 | 0 | 85 | **0.0** | | 25% of cells show SFPQ localised to nuclear envelope |
| UT 1 | | | | | | 0 | 122 | **0.0** | | |
| UT 2 | | | | | | 0 | 85 | **0.0** | | |
